# Recruitment of CTCF to an *Fto* enhancer is responsible for transgenerational inheritance of obesity

**DOI:** 10.1101/2020.11.20.391672

**Authors:** Yoon Hee Jung, Hsiao-Lin V. Wang, Daniel Ruiz, Fiorella C. Grandi, Brianna J. Bixler, Hannah Linsenbaum, Jian-Feng Xiang, Samantha Forestier, Andrew M. Shafik, Peng Jin, M. Ryan Corces, Victor G. Corces

## Abstract

Transgenerational transmission of epiphenotypes is poorly understood. Here we show that exposure of pregnant mouse F0 females to BPA results in obesity in the F2 progeny due to increased food intake and leptin resistance. This epiphenotype can be transmitted up to the F6 generation and disappears in F7. Analyses of chromatin accessibility in sperm of the F1-F6 generations reveals alterations in the binding of CTCF at two enhancers of the *Fto* gene in obese but not control animals that correlates with transmission of obesity. Deletion of the CTCF site in *Fto* results in mice that fail to become obese when exposed to BPA. These *Fto* enhancers show increased interactions in sperm of obese mice with the *Irx3* and *Irx5* genes, which are involved in the differentiation of appetite controlling AgRP/NPY neurons. Single-nucleus and immunofluorescence analyses in the arcuate nucleus of the hypothalamus suggest that exposure to BPA results in expansion of the number of orexigenic AgRP neurons. This expansion correlates with increased accessibility of the *Fto* proximal enhancer in radial glia-like neural stem cells (RG-NSCs), which give rise to AgRP/NPY neurons, and in mature oligodendrocytes. The results provide a molecular mechanism for transgenerational inheritance in mammals and suggest that both genetic and epigenetic alterations of *Fto* can lead to the same phenotypic outcomes.

## Introduction

Evidence suggestive of inter and transgenerational inheritance of epiphenotypes in mammals has been reported extensively. Numerous studies suggest a link between parental environments and a variety of effects in the offspring (*1–3*). For example, exposure of laboratory animals or humans to endocrine disrupting compounds leads to increased reproductive dysfunction, cancer, obesity, diabetes, and behavioral disorders (*4*). However, we still lack an understanding of the molecular processes by which epigenetic alterations induced by environmental exposures are transmitted between the exposed and subsequent generations in mammals. This is in part due to the peculiar state of the nucleus in the mature gametes, which is highly condensed in sperm and arrested in metaphase II in mature oocytes. In addition, the epigenetic content of the mammalian germline in the form of DNA methylation is reprogrammed at least twice in each life cycle, once during the differentiation of the primordial germ cells (PGCs) and a second time after fertilization during pre-implantation development of the embryo (*5*). Histone modifications are also reprogrammed during PGC development. Mouse and human sperm retain 8-15% of the histones present in somatic cells, and these histones contain covalent modifications characteristic of active and silenced chromatin at specific genes (*6–11*). Many of these histone modifications are erased after fertilization when protamines are replaced by histones in the zygote, bringing into question whether histone modifications can carry epigenetic information between generations (*12*). Mechanisms involving small RNAs have been successful in explaining transgenerational phenomena in *C. elegans*, where these RNAs can be amplified by RNA-dependent RNA polymerases in each generation (*13*). The contribution of small RNAs present in the gametes to the transmission of epiphenotypes between generations has also been explored in mammals, perhaps in part due to the apparent lack of reprograming-resistant epigenetic information that can serve as memory of ancestral environmental influences. For example, tRNA fragments made in the epididymis and transported to sperm have been implicated in intergenerational transmission of metabolic effects caused by paternal protein restriction or high fat diet (*14, 15*). miRNAs present in sperm and transferred to the zygote may cause changes in gene expression affecting the differentiation of adult tissues, and have been suggested to be responsible for the transmission of behavioral and metabolic changes caused by early life traumatic stress (*14–17*). However, there is no current evidence suggesting that small RNAs are able to transmit environmentally-induced phenotypes through multiple generations in mammals.

Transcription factors (TFs) sit at the top of the cascade of events that lead to gene expression and are causal to many of the chromatin features normally considered to carry epigenetic information. It is thus possible that TF occupancy cooperates with DNA methylation or covalent histone modifications to retain epigenetic information through reprogramming events in the germline or the early embryo. This has been shown to be the case for TFs that protect their binding sites from re-methylation both during germline development and in the blastocyst (*18*). A variation of this idea is represented by the interplay between placeholder nucleosomes containing the histone variant H2A.Z and DNA methylation, which has been shown to mediate transmission of information between the gametes and embryos in zebrafish (*19*). Therefore, memory of ancestral epigenetic alterations caused by environmental changes may be encoded by a combination of reprogramming-resistant molecular events during critical periods of the life cycle, such as germline differentiation and pre-epiblast development, together with more classical epigenetic information at other stages of cell differentiation.

Here we use bisphenol A (BPA) to induce an obesity epiphenotype that can be transmitted down to the F6 generation through the male and female germlines. Using the assay for transposase-accessible chromatin by sequencing (ATAC-seq) to identify sites in the genome of the gametes bound by TFs and correlating this information with transmission of obesity, we identify several sites containing motifs for CTCF, FOXA1, Estrogen receptor alpha (ESR1) and Androgen receptor (AR) present in two enhancers of the *Fto* gene in obese but not control mice. Mouse strains carrying a deletion of a 300 bp region containing the CTCF and FOXA1 sites in an enhancer of *Fto* fail to transmit the obesity epiphenotype to subsequent generations when exposed ancestrally to BPA. Finally, we trace the reasons for obesity to an increase of orexigenic AgRP neurons in the arcuate nucleus of the hypothalamus of BPA mice.

## Results

### Exposure of pregnant females to BPA causes obesity that can be transmitted transgenerationally

To gain insights into the mechanisms by which epiphenotypes elicited by environmental exposures can be transmitted transgenerationally, we examined the effect of the endocrine disrupting chemical BPA. Previous work has shown that exposure of pregnant rats and mice to a variety of endocrine disruptors during the window of time when the germline of the embryo is demethylated leads to various health effects in the progeny, including testicular and breast cancer, behavioral defects, and obesity (*20, 21*). Following this work, we exposed two pregnant female mice (F0) via daily intraperitoneal injection of BPA (50 mg/kg body weight/day) during days E7.5 through E13.5 of fetal development (*22*). As a control, two pregnant females were injected with vehicle only (see Methods section for details). In the rest of the manuscript, we will refer to the progeny of BPA-exposed F0 females as BPA-Fi, where i is the generation number, and the progeny of vehicle-exposed females as CTL-Fi, although the exposure took place only in the F0 generation. The BPA-F1 progeny has no overt phenotypic abnormalities when compared to CTL-F1 and their body weight is the same as that of controls (Figure 1A). BPA-F1 males were then crossed to BPA-F1 females from a parallel exposure experiment (Figure S1A). BPA-F2 males and females displayed an obvious increase in body weight with respect to CTL-F2 animals (Figure 1B). It has been estimated that humans are exposed to an average of 50 mg/kg body weight of BPA (*23*), resulting in concentrations of this compound in serum in the ng per ml range (*24*). Since the amount of BPA to which mouse pregnant females are exposed in our experiments is much higher than that to which the average human is thought to be exposed, we injected pregnant females with decreasing concentrations of BPA and examined the effect on body weight in the progeny. The results indicate that BPA at concentrations of 50 μg/kg, which are comparable to those to which humans are normally exposed, cause a similar increase in body weight as higher concentrations (Figure S1B). To determine the amount of BPA in the exposed embryo compared to that injected in pregnant females, we measured BPA levels using mass spectroscopy at different times after injection. We found that the amount of BPA in embryonic tissues is 1.5 mg/kg 10 min after injection, rapidly decreases by 500-fold to 0.105 mg/kg after 120 min and is undetectable by 24 h (data not shown). This suggests that BPA is rapidly eliminated or metabolized and that a very transient daily exposure is able to elicit the effects described here.

**Figure 1.**
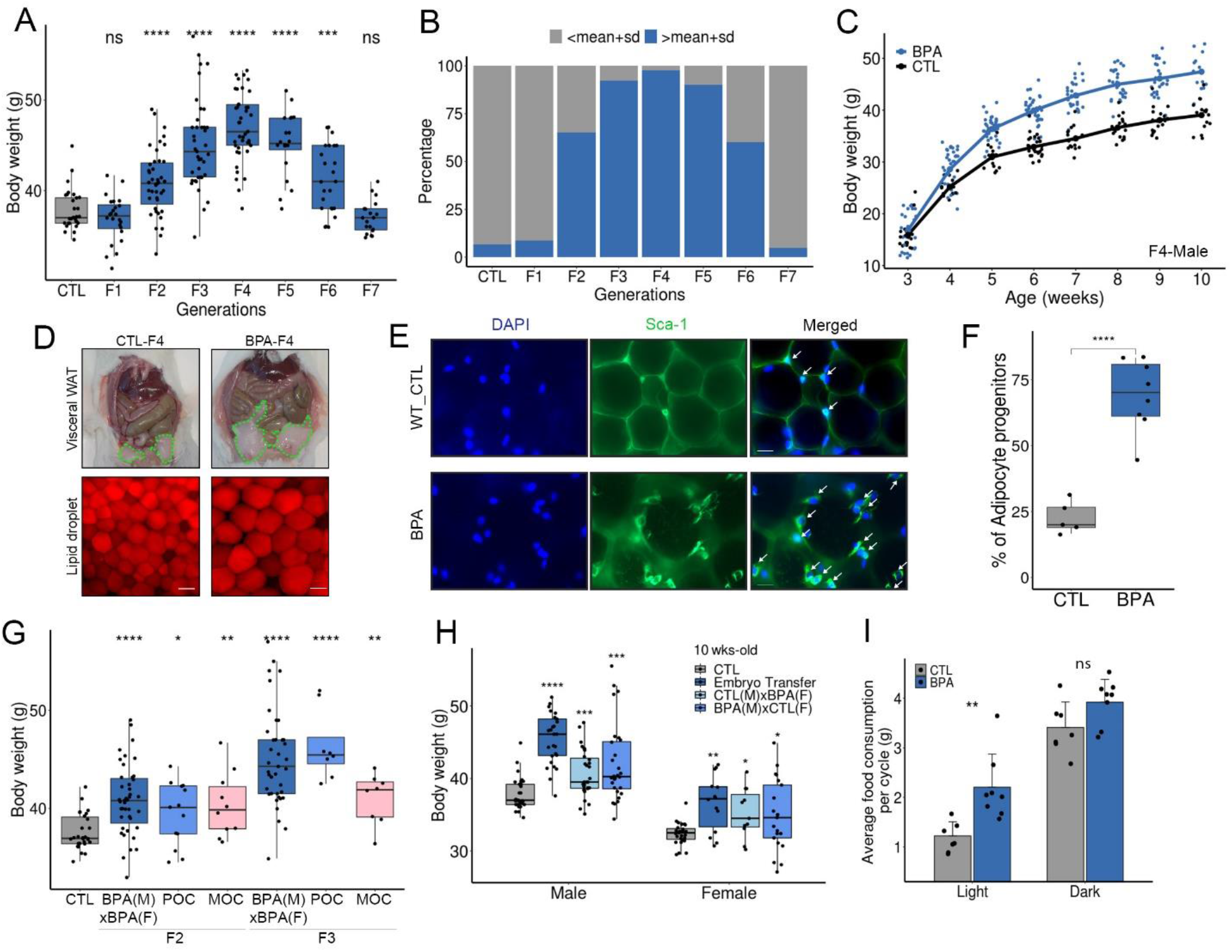
Analysis of BPA-induced obesity. (A) Changes in body weight between control and different generations of the progeny of BPA-exposed F0 pregnant female mice (CTL n=31, F1 n=24, F2 n=43, F3 n=39, F4 n=41, F5 n=20, F6 n=25, F7 n=19). (B) Percentage of overweight mice in the F1-F7 generation progeny of F0 BPA-exposed pregnant females. (C) Changes in body weight with age in control and the F4 generation progeny of BPA-exposed females (CTL n=17, BPA n=26). (D) Differences in visceral fat, stored lipids, and adipocyte size in BPA-F4 versus control; bar represents 50 μm. (E) Immunofluorescence microscopy of visceral fat tissue from control and BPA-F4 mice stained with antibodies to Sca-1, which labels adipocyte progenitor cells. (F) Fraction of adipocyte progenitor cells present in the visceral fat tissue of control and BPA-F4 mice (CTL n=5, BPA n=8). (G) Differences in body weight in F2 and F3 generation male mice ancestrally exposed to BPA and outcrossed to controls. POC, paternal outcross in which a BPA female was crossed to a control unexposed male. MOC, maternal outcross in which a BPA male was crossed to a control unexposed female (CTL n=30, F2; BPA (M) x BPA (F) n=43, MOC n=10, POC n=13, F3; BPA (M) x BPA (F) n=39, MOC n=8, POC n=8. (H) Body weight differences in males and females between control, progeny from embryo transfer (ET) experiments, and outcrosses between BPA-F2 and control (Male; CTL n=31, Embryo Transfer n=27, CTL (M) x BPA (F) n=31, BPA (M) x CTL (F) n=28, Female; CTL n=27, Embryo Transfer n=14, CTL (M) x BPA (F) n=11, BPA (M) x CTL (F) n=21. (I) Average food intake by BPA-F4 and CTL-F4 males during the light and dark cycles over a 3-day period (CTL n=7, BPA n=8). For all panels, P values were calculated using t-test with respect to control; **** p<0.0001, ***p<0.001, ** p<0.01, *p<0.05, ns not significant.

Crosses between BPA-Fi males and females selected to have a weight equal to the median for each generation (Figure 1A) were continued as described in Figure S1A. The increased body weight was also observed in the BPA-F3 through BPA-F6 generations, but it was lost in BPA-F7 animals (Figure 1A). To compare changes in body weight in different generations, we took the median weight of control males at week 10 and we defined obese animals as those whose weight is one standard deviation above the median control weight. Only 5% of males are overweight in the BPA-F1 generation by this criterion, which is similar to control (Figure 1B). The fraction of overweight males increases to 65% in BPA-F2, peaks at 97% in BPA-F4, and starts decreasing through BPA-F5-6. Approximately 60% of F6 males are obese and 40% have normal weight; we will refer to these F6 mice with normal weight as “lean”. All BPA-F7 males display weights similar to those of control males (Figure 1B). Body weight of BPA-Fi mice of any generation is the same as that of controls at 3 weeks of age, but differences become statistically significant by weeks 5-6 and remain for the rest of adult life (Figure 1C). These differences are more pronounced in males than females (Figures 1C and S1C). Crosses between overweight or between lean BPA-F6 animals produce progeny with weights statistically indistinguishable from those of controls (Figure S1D), suggesting that the germline of F6 animals has lost the information responsible for the inheritance of obesity independent of the weight of the animals. The median weight for CTL-F4 males is 39.2 g and for BPA-F4 males is 51.1 g (Figure S1E). BPA-Fi mice show a large accumulation of visceral white adipose tissue (WAT) compared to controls (Figure 1D, top), and increased number of adipocytes and enlargement of lipid droplets (Figure 1D, bottom). Whereas the visceral WAT represents 1.32% of total body weight in control male animals, this value increases to 2.45% in BPA-F4 males ancestrally exposed to BPA (Figure S1F). To understand the origin of the higher number of adipocytes present in BPA-F4 fat tissue, we examined its composition in more detail by performing immunofluorescence microscopy with antibodies to stem cell antigen-1 (Sca-1), which labels adipocyte progenitor cells (Figure 1E). Results show that 69% of the cells in fat tissue from BPA-F4 males correspond to adipocyte progenitors compared to 22% in the fat tissue of control animals (Figure 1F).

### BPA-induced obesity is a result of increased food consumption

In the experiments described above, obesity was observed in the F2 progeny from two F1 mice from two parallel different exposures (Figure S1A). Therefore, BPA could have affected either of the two parental germlines or both. To determine whether the obese epiphenotype can be inherited through the paternal or maternal germlines, we outcrossed the F1 progeny to control unexposed mice of the opposite sex (Figure S2A). We then measured the body weight of males from the F2 and F3 generations. We find that the median weight increases in the F2 progeny and raises further in the F3 independent of the parental origin of the exposed germline, suggesting that the obese epiphenotype can be inherited through both the paternal and maternal germlines (Figure 1G).

To ensure that the obese phenotype is not the result of external factors such as the maternal environment, transfer of the maternal microbiome, presence of constituents other than sperm in the seminal fluid, etc., we performed *in vitro* fertilization using purified oocytes from a BPA-F2 female and sperm isolated from the cauda epididymis of a CTL-F2 male. The resulting embryos were then implanted into pseudo pregnant unexposed control females and the weight of their progeny was measured at 10 weeks of age. As controls, CTL-F2 males were directly crossed with BPA-F2 females and vice-versa. The results indicate that the median weight of males and females from the embryo transfer experiment is significantly higher than that of unexposed controls (Figure 1H), suggesting that the overweight phenotype is indeed due to epigenetic changes in the germline rather than external factors.

To gain insights into the physiological processes underlying the obesity phenotype, we housed mice in metabolic cages and measured food intake and energy use. BPA-F4 mice show an increase in total food consumption with respect to controls (Fig. S2B). This increase is due to additional eating during the fasting light cycles (Figure 1I). In agreement with this observation, control mice show a lower respiratory exchange ratio during the fasting light cycle than during the active dark cycle, indicative of preferential fat utilization during the day when they are asleep (*25*). BPA-F4 mice show similar respiratory exchange ratios during the light and dark cycles indicative of constant carbohydrate preference throughout the 3-day metabolic cage test (Figure S2C). However, the energy expenditure of control and BPA-F4 mice is the same (Figure S2D). Similar results were obtained by comparing overweight and lean BPA-F6 mice. BPA-F6 overweight mice consume more food (Figure S2E), with increased consumption during the light cycle (Figure S2F), but they spend the same energy as their lean siblings (Figure S2G). Together, these results suggest that obese mice ancestrally exposed to BPA exhibit unregulated eating patterns that reduce their ability to utilize fat as a fuel source, and expend the same energy as control mice, resulting in increased adiposity.

### Exposure to BPA results in alterations of CTCF occupancy and 3D interactions in the germline

The results described so far underscore two important questions i.e., how does the obesity epiphenotype arise and what are the mechanisms by which this phenotype is transmitted transgenerationally. We will address the latter question first because this information is important in explaining the obesity epiphenotype. Since BPA is an estrogen-like compound that has been shown to act through a variety of nuclear hormone receptors (*26*), we hypothesized that exposure to BPA may cause alterations in transcription accompanied by a redistribution of TFs throughout the genome of the germline during E7.5-E13.5, the time of embryonic development when the DNA of germline cells becomes demethylated. The unmethylated state of the DNA may allow binding of TFs to new sites in the genome where these TFs cannot bind when the DNA is methylated. Subsequently, the presence of bound TFs at these new sites may protect the DNA from re-methylation and these changes may persist in the mature germline (*18*). The transgenerational transmission of obesity decreases substantially in the F6 generation, suggesting that epigenetic alterations induced by exposure to BPA start disappearing in the sperm of F5. Therefore, to uncover alterations in TF binding responsible for transgenerational inheritance, we performed ATAC-seq in sperm isolated from the cauda epididymis of control and BPA F1-F6 males. We used sub-nucleosome-size reads, which correspond to the presence of bound TFs in chromatin (*27*) and we will refer to these as Transposase Hypersensitive Sites (THSSs). We first identified THSSs present in generations BPA-F1-F4 with respect to control using an 0.05 FDR and ≥ 2-fold change cutoff and requiring that differences are consistently present in all replicates of each sample. Using these criteria, we identified 69 sites that are present in two different replicates of sperm from BPA F1-F4 males but not in controls (Figure 2A and Figure S3A). Most of the 69 BPA-gained sites are present in distal intergenic regions or introns, suggesting that they may correspond to regulatory sequences such as enhancers (Figure S3B). Motif analysis at the summits of ATAC-seq peaks at these sites indicates the presence of binding motifs for a variety of TFs such as CTCF, ZNF143, FOXA1, and several nuclear hormone receptors, including ESR1/2, AR, and PPARγ (Figure 2B). An example of differential ATAC-seq peaks containing binding sites for CTCF and AR upstream of the *Tram2* gene is shown in Figure S3C. Thirty-four of these sites contain binding motifs for CTCF plus another TF within the same ATAC-seq peak (Figure S3D). In addition to the 69 ATAC-seq peaks gained in the sperm of mice ancestrally exposed to BPA but not in controls, we also identified 5 lost sites (Figure S3A). Visual inspection of these 5 sites suggests that they correspond to peaks with very low ATAC-seq signal in sperm of control males (Figure S3E). Since their biological significance is unclear, they will not be considered further.

**Figure 2.**
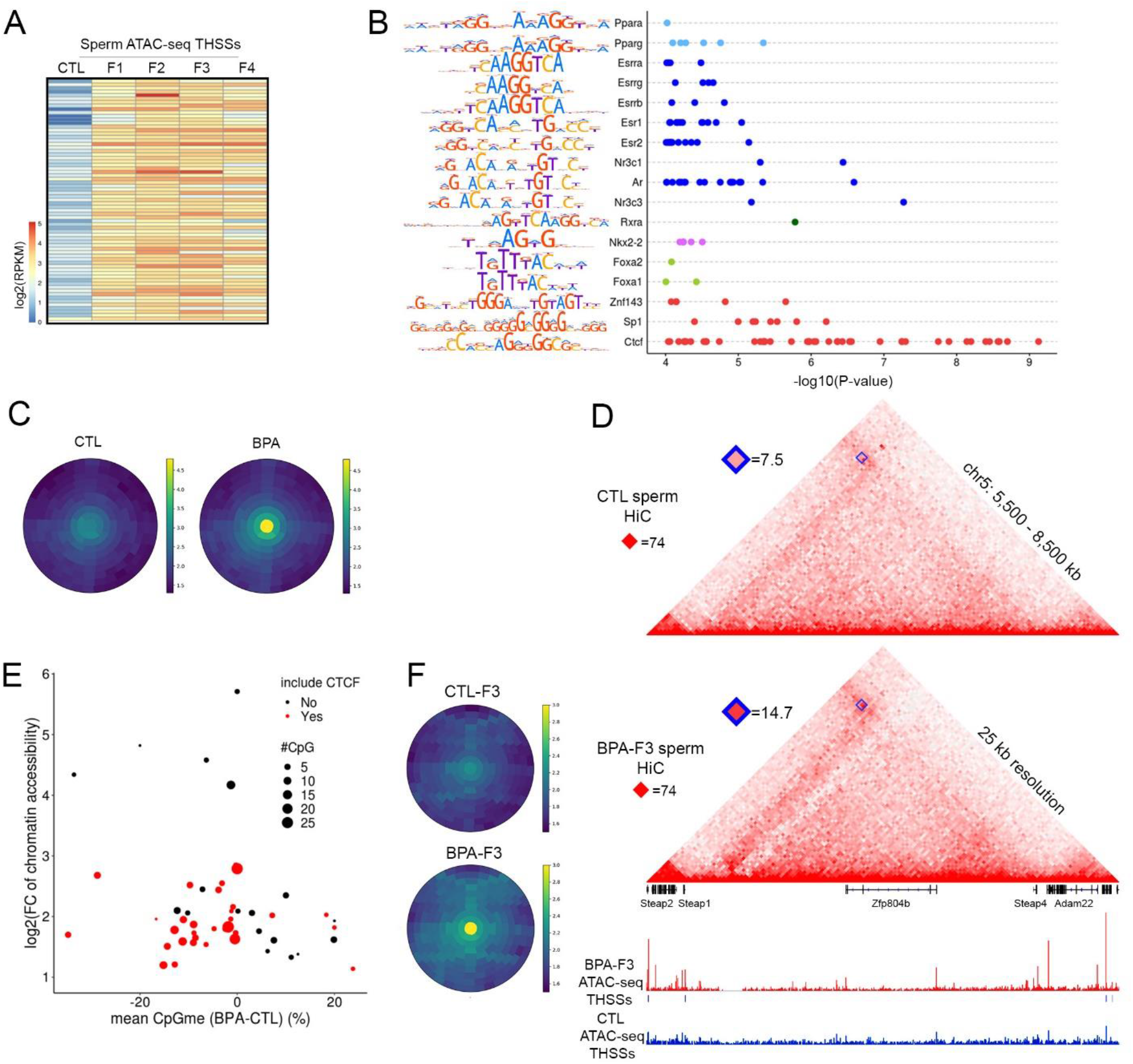
Effect of BPA on TF occupancy and 3D genome architecture. (A) Heatmap of ATAC-seq sites differentially accessible in sperm between CTL-F1 and BPA-F1-F4. (B) TFs whose binding motifs are enriched at ATAC-seq accessible sites present in sperm of BPA-F1 through BPA-F4. (C) Frequency of interactions mediated by CTCF sites differentially accessible in sperm of CTL-F1 versus BPA-F1-F4; Hi-C data is from sperm of F3 males. (D) Specific example of a CTCF loop whose anchors interact more frequently in sperm of BPA-F3 than CTL-F3. (E) Differences in DNA methylation in sperm between BPA-F3 and CTL-F3 at ATAC-seq sites conserved in BPA-F1 through BPA-F4. (F) Interaction frequencies between ATAC-seq sites conserved in sperm of BPA-F1 through BPA-F4 and other sites in the genome calculated using significant interactions from Hi-C data obtained in sperm of BPA-F3 and CTL-F3.

To test whether the new CTCF sites observed in sperm of BPA mice affect the 3D organization of sperm chromatin, we performed Hi-C (see Supplemental Table 1 for quality controls of the Hi-C libraries) in sperm of BPA-F3 and CTL-F3, and compared interactions between anchors containing differential CTCF sites and other CTCF sites in the genome using SIP (*28*). Results indicate a dramatic increase in interactions observed in Hi-C heatmaps as punctate signal, indicating the formation of new CTCF loops (Figure 2C). An example of changes in CTCF levels in BPA-F3 sperm and the formation of a loop between CTCF sites adjacent to *Steap1* and *Steap4*, two genes shown to be involved in obesity in humans (*29*), is shown in Figure 2D. These results suggest that BPA, directly or indirectly, induces the binding of CTCF and other TFs at different regions in the genome in the sperm of the F1 generation, many of which are conserved in subsequent generations, altering the 3D organization of chromatin in mature sperm.

### Sites with altered occupancy of TFs in sperm are differentially methylated

To explore the relationship between the presence of these TFs at new sites in the sperm genome and possible changes in DNA methylation that could account for their differential binding to DNA, we performed genome wide bisulfite sequencing (GWBS) in sperm of control and BPA-F3 males. We find a large number of differentially methylated regions (DMRs) defined as a change of >20% in methylation levels and a p value ≤0.0001 (Figure S3F). Of these, 1,428 regions are hypomethylated in sperm of BPA-F3 with respect to control and 648 are hypermethylated, suggesting that exposure of the germline to BPA induces DMRs that persist transgenerationally. These DMRs are located almost exclusively in distal intergenic and intronic regions, suggesting that they may correspond to enhancers (Figure S3G). We then explored the relationship between the observed changes in ATAC-seq signal between BPA and control versus the change in CpG methylation in a 300 bp region surrounding the summit of the 69 ATAC-seq peaks maintained in BPA-F1 through BPA-F4. Sites with increased accessibility in BPA sperm are generally hypomethylated with respect to control, especially those containing the binding motif for CTCF, whose interaction with DNA is known to be sensitive to methylation (Fig. 2E and Figure S3D bottom tracks). These observations agree with the possibility that BPA-induced alterations of transcription in PGCs at the time when the DNA is being demethylated may allow binding of CTCF and other TFs to specific sites in the genome, which are then protected from re-methylation when this process occurs after E13.5 in the male germline. These sites may then become enhancers in an epigenetically active state in sperm. To explore this possibility, we called significant interactions in the Hi-C data using Fit-Hi-C (*30*) and identified genes whose promoters are contacted by the 69 putative enhancers in BPA-F3 sperm. Results show that a total of 610 gene promoters are contacted by these 69 sites. Metaplots of interaction frequencies show an increase in contacts between the putative enhancers and the promoters of target genes in BPA-F3 versus control sperm (Figure 2F). Of the 610 genes contacted by the 69 BPA-induced differential sites, 30 are involved in obesity based on GWAS studies performed in humans. Furthermore, mouse carrying mutations in genes contacted by the 69 sites with increased accessibility show defects in beta cell physiology, hepatic and adipose tissue morphology, and hormone secretion (Figure S3H). Together, these results suggest that BPA, directly or indirectly, induces the binding of CTCF and other transcription factors at genomic sites that are normally methylated. Many of these sites become hypomethylated with respect to control in the sperm of BPA animals. It is possible that these sites correspond to enhancers that become activated in the BPA male germline. These activated enhancers contact gene promoters at a higher frequency in BPA sperm. As we have previously shown, this information may be transmitted to the embryo after fertilization (*12*), resulting in alterations in transcription during early embryogenesis.

### Activation of an enhancer in the *Fto* gene in the male and female gametes correlates with transmission of obesity

Since the obesity epiphenotype starts decreasing in the F5 generation, is limited to approximately 60% of the progeny in F6, and it completely disappears in F7, we examined the correlation between the transmission of the phenotype and the presence of the 69 altered TF sites in sperm from these later generations. Only 12 out of the 69 sites are absent in sperm of F6 compared to F4, suggesting that all or some of these 12 sites, but probably not the rest of the 69, are responsible for the transgenerational transmission of obesity. One of these 12 sites is present in intron 8 of the *Fto* gene (Site 1, Figure 3A), which encodes a demethylase of N6-methyladenosine (m^6^A) in RNA. SNPs in the first two introns of this gene in humans have been implicated in obesity based on results from GWAS studies (*31*). Site 1 in intron 8 of *Fto* contains motifs for the CTCF and FOXA1 proteins. ChIP-seq experiments confirm the presence of CTCF at site 1 in the *Fto* gene in BPA-F3 but not control mice (Figure 3A, lower panel). Adjacent to this site, several additional ATAC-seq peaks containing motifs for ESR1, AR, and PPARγ are either only present in sperm of BPAF1-F5 or their intensity increases in sperm from these animals with respect to BPA-F6 and controls (Sites 2-6, Figure 3A). These sites were not identified in our original analysis for differential ATAC-seq peaks between BPA-Fi and control because of the stringent statistical conditions imposed in the analysis; however, visual inspection clearly indicates the existence of differential accessibility in a large region of the *Fto* intron containing Sites 1-6 (Figure 3A). These same sites are also accessible in the sperm of male progeny obtained from the embryo transfer experiments (Figure 3A). We will refer to this approximately 4 kb region in intron 8 of the *Fto* gene whose accessibility increases in sperm of the male progeny of BPA-exposed females as the “*Fto* BPA proximal enhancer”.

**Figure 3.**
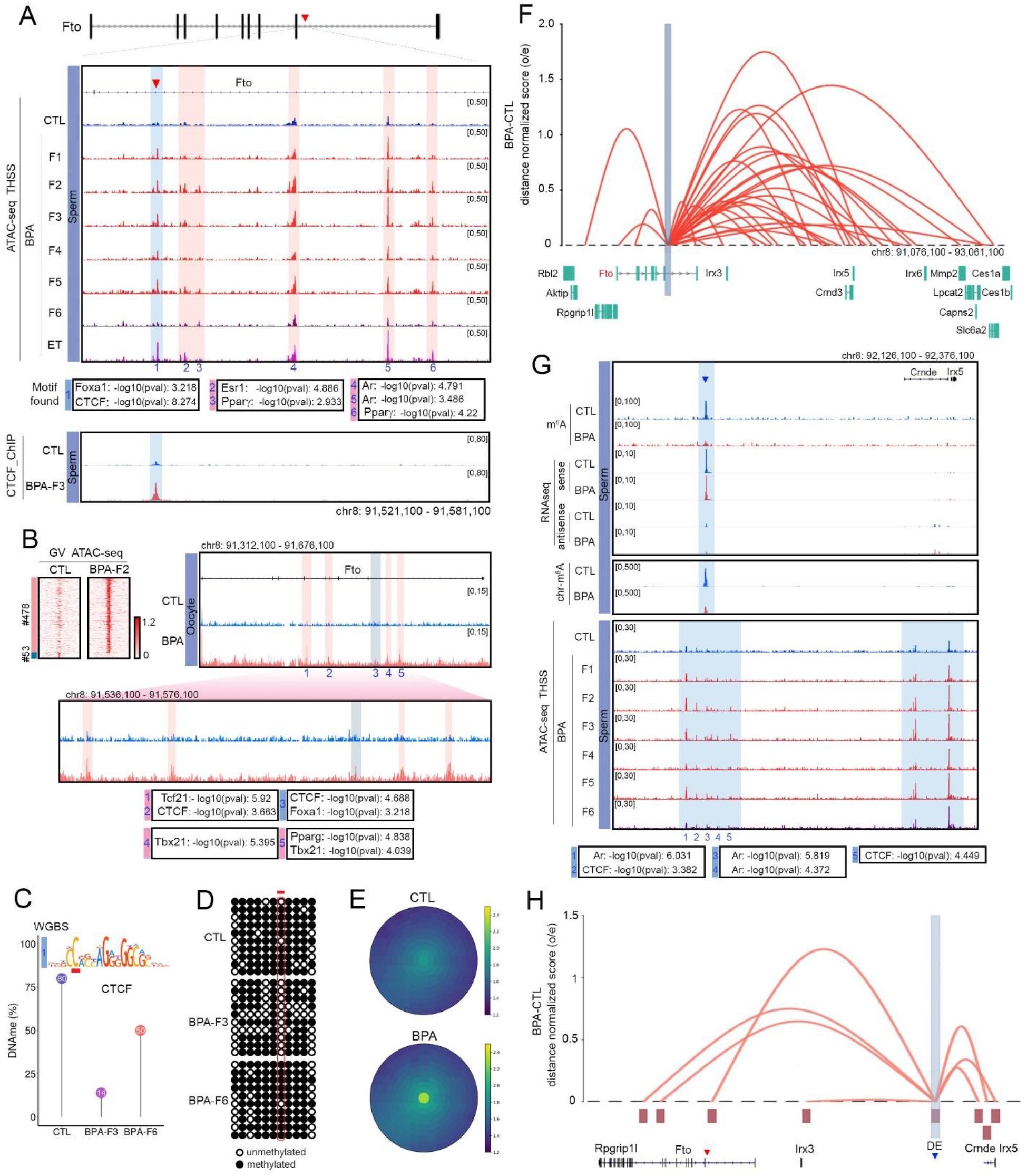
Changes in accessibility and interactions at an *Fto* intronic enhancer correlate with transmission of obesity. (A) Persistence of an ATAC-seq accessible site (Site 1) located in intron 8 of the *Fto* gene in sperm of BPA-F1 through BPA-F5 males. This site contains motifs for CTCF and FOXA1, and it is also present in the sperm of the male progeny from the embryo transfer (ET) experiment. Other sites in the region are present in CTL and BPA-F6 but they increase in signal in sperm of BPA-F1 through BPA-F5 (top). ChIP-seq experiments confirm the presence of CTCF at site 1 in sperm of BPA-F3 but not control males (bottom). (B) ATAC-seq sites differentially accessible between oocytes of BPA-F2 and CTL-F2 females. A subset of these sites is located in several introns of *Fto* and one of them coincides with Site 1 present in intron 8 in sperm. (C) Methylation levels at the CpG located in position 2 of the CTCF motif present in Site 1 obtained from WGBS-seq in sperm of CTL, BPA-F3 and BPA-F6. (D) Methylation levels at the same CpG obtained from bisulfite sequencing. (E) Differences in interaction frequency between the *Fto* proximal enhancer and other sites in the genome using significant interactions obtained from Hi-C data in sperm of CTL-F3 and BPA-F3 mice. (F) Differential Hi-C interactions between the *Fto* proximal enhancer and adjacent genomic sites in sperm of BPA-F3 and CTL-F3 mice. (G) RNA-seq and m^6^A levels in an enhancer RNA present distally from *Fto* in a region close to the *Crnde* and *Irx5* genes. This putative enhancer contains binding sites for several TFs whose accessibility increases in BPA-Fi with respect to CTL and BPA-F6. (H) The *Fto* distal enhancer preferentially interacts with the *Fto* and *Irx5* genes in sperm of BPA-F3 versus CTL-F3.

Since the obesity epiphenotype elicited by BPA exposure can also be transmitted through the female germline, alterations in TF binding responsible for obesity should also be present in oocytes. To explore this possibility, we performed ATAC-seq in oocytes from control and BPA-F2 females. Results show that GV oocytes from BPA-F2 females have 478 accessible sites and lack 53 sites with respect to CTL-F2 oocytes (Figure 3B). Out of the 12 sites present in BPA-F1-F5 but lost in BPA-F6 sperm, only Site 1 present in intron 8 of the *Fto* gene in sperm of BPA-Fi is also differentially accessible in oocytes (Figure 3B), suggesting that *Fto* may be the primary effector of transgenerational inheritance of obesity whereas other differentially accessible sites in the genome may be secondary consequences of *Fto* activation or unrelated to the obesity phenotype. Sites 2-6 present in intron 8 of *Fto* in sperm of BPA-Fi males are not present in oocytes of BPA-F2 females. Instead, different sites in introns 4, 5, and 8 are differentially accessible in oocytes of BPA-F2 females with respect to controls. These sites contain binding motifs for CTCF, FOXA1, TCF21, TBX21 and PPARγ. One interesting consequence of results obtained in oocytes of BPA-F2 females and those obtained in sperm after the embryo transfer experiments using the same BPA-F2 oocytes is that the transcription factor sites present in the male gametes of the embryo transfer progeny have switched from the oocyte to the sperm distribution patterns (Figures 3A-B).

Since binding of CTCF to DNA is known to be sensitive to the methylation status of a specific C located at position 2 in the core motif (*32*), we examined base pair resolution GWBS data obtained in sperm from control, F3, and F6 mice for changes in DNA methylation at the CpG located in this position. Results show that the level of methylation in this CpG is around 80% in sperm from control males, it decreases to 14% in sperm from BPA-F3 and increases to 50% in BPA-F6 sperm (Figure 3C). To determine the methylation levels of the CpG present at position 2 in the CTCF motif in Site 1 more quantitatively, we performed bisulfite conversion followed by Sanger sequencing. We find that this C is highly methylated in sperm from control males, is completely demethylated in BPA-F3 sperm and becomes almost completely remethylated at levels similar to control in BPA-F6 sperm (Figure 3D). These results again agree with the idea that binding of TFs to the unmethylated DNA of the germline may protect from DNA remethylation to serve as the memory underlying the transgenerational transmission of environmentally-induced epiphenotypes.

BPA-induced occupancy by TFs of the *Fto* BPA proximal enhancer may result in increased interactions with its target genes. To test this possibility, we analyzed Hi-C data obtained in sperm of control and BPA-F3 mice using significant interactions identified by Fit-Hi-C (*30*). The results show an increase in interaction frequency in sperm of BPA-F3 mice between the *Fto* proximal enhancer and adjacent sites in the genome, including its own promoter as well as neighboring genes (Figures 3E and 3F). Several of these genes, including *Rpgrip1l*, *Irx3*, *Irx5*, *Slc6a2*, and *Mmp2*, have been shown to be involved in the regulation of body size and obesity by affecting appetite and food consumption (*33–37*), which agrees with observations indicating that BPA-F4 mice have increased food consumption (Figure 1I). However, the transcript levels for these genes, except for Mmp2, are not statistically significantly different between control and BPA sperm (Figure S4A). Since sperm are transcriptionally inactive, it is possible that the increased interactions between the *Fto* BPA proximal enhancer and target promoters are a vestige of previous transcription during germline development and may have a functional effect after fertilization during pre-implantation development (*12*).

### m^6^A demethylation of a distal enhancer RNA may also affect *Fto* expression

Since *Fto* encodes an m^6^A demethylase, we tested whether altered expression of this gene during spermatogenesis affects RNA methylation by measuring m^6^A levels in sperm RNAs of BPA-F3 and controls. We find a large global decrease in this modification in RNA from sperm of BPA-F3 males with respect to control (Figure S4B). The absence of m^6^A does not appear to affect stability of sperm RNAs, since global levels of both sense and antisense RNAs are similar in BPA-F3 and control (Figure S4C). Although some of the m^6^A hypomethylated sites map to exons of mRNAs (Figure S4D), most are located in introns and intergenic regions, suggesting that they may be present in enhancer RNAs (Figure S4E) It is possible that the low levels of m^6^A in sperm RNAs of BPA males will alter their function during early embryonic development after being transferred to the zygote.

It has been reported recently that m^6^A hypomethylated eRNAs preferentially localize to chromatin where they increase accessibility (*38*). Analysis of Hi-C data using Fit-Hi-C indicates that putative enhancers in which m^6^A levels decrease significantly in sperm of BPA-F3 with respect to control interact with promoters of genes involved in various processes including development and response to stimuli (Figure S4F). We thus examined the region surrounding the *Fto* locus for putative eRNAs. We noticed the expression of a ncRNA in the region between the *Irx3* and *Irx5* genes (Figure 3G, top panel; see Figure 3F for the relative location of these two genes with respect to *Fto*). Although the levels of this RNA remain the same, the RNA is m^6^A hypomethylated in BPA-F3 sperm with respect to control (Figure 3G, top panel). To further explore the significance of this putative eRNA, we isolated chromosome-associated RNAs from BPA-F3 and control sperm. We then performed RNA-seq and determined m^6^A levels in chromatin-associated RNAs. Results indicate that the putative eRNA is tightly bound to chromatin and the chromatin-associated fraction is hypomethylated in BPA-F3 sperm (Figure 3G). Interestingly, this RNA is transcribed from a region that is not accessible in control mice, but it becomes accessible in BPA-F1-F5 sperm and loses accessibility in sperm from the F6 generation (Figure 3G, lower panel). This region was not identified in our initial analyses because the low ATAC-seq signal in the sperm of the F4 generation did not meet the established statistical threshold criteria. The main ATAC-seq peak in the region contains motifs for CTCF and AR. Other adjacent sites also contain motifs for these two proteins and thus this region could represent an enhancer. Analysis of Hi-C data from BPA-F3 and control sperm using Fit-Hi-C indicates that this putative enhancer preferentially interacts in BPA with respect to control with the promoter of the *Fto*, *Irx3*, and *Irx5* genes, as well as the *Fto* BPA proximal enhancer identified above (Figure 3H), suggesting that it may regulate the expression of these genes. We will refer to this enhancer as the “*Fto* BPA distal enhancer”.

### Mice carrying a deletion of the FOXA1/CTCF sites do not become obese after ancestral BPA exposure

Transmission of BPA-induced obesity correlates with increased occupancy of CTCF and several TFs at the *Fto* BPA proximal and distal enhancers. To test the involvement of these enhancers in the transgenerational transmission of obesity more directly, we used CRISPR to delete a small region surrounding the FOXA1 and CTCF sites in the *Fto* proximal enhancer. The binding sites for these two proteins are separated by approximately 90 bp and we obtained two different mutant strains carrying deletions of 343 bp and 349 bp sequences (Figure S5A), which we will refer to as *Fto^c1^* and *Fto^c2^*. Neither of these two strains displays any obvious observable phenotypes. After outcrossing, we obtained homozygous strains and examined their response to BPA exposure. The results of these experiments were indistinguishable between the *Fto^c1^* and *Fto^c2^* mutant strains and the data presented here as *Fto^c^* is the combination of results obtained for each strain separately. *Fto^c^* females were exposed to BPA (*Fto^c^*_BPA) or vehicle (*Fto^c^*_CTL) and F1 progeny from independent exposures were crossed to obtain the F2 generation. Contrary to wild type mice (Figure 1A), *Fto^c^* mice failed to become obese after ancestral exposure to BPA (Figures S5B and S5C). F2 animals were then crossed as described above (Figure S1A) and the F3 progeny was analyzed in detail. F3 *Fto^c^*_BPA males failed to gain excess weight with age and their weight was indistinguishable from that of *Fto^c^*_CTL or WT_CTL males (Figure 4A). F3 *Fto^c^*_BPA males show no increase in visceral fat (Figures 4B and S5D) and the size of the adipocytes and amount of lipid droplets is normal (Figure 4B, lower panel). We also examined the adipocyte progenitor cells present in the visceral fat tissue of *Fto^c^* males by immunostaining with SCA-1 antibodies (Figure S5E). The relative number of adipocyte progenitors does not increase as a consequence of BPA exposure, and it remains at the same levels in *Fto^c^*_BPA and *Fto^c^*_CTL as in unexposed WT controls (Figure 4C). F3 *Fto^c^* males ancestrally exposed to BPA have a similar food intake as unexposed mice (Figure S5F), and they consume the same amount of food as controls during the light and dark cycles (Figure 4D). Furthermore, neither the respiratory exchange ratio nor energy expenditure are affected by ancestral BPA exposure in mice carrying the deletion of the FOXA1/CTCF sites (Figure 4E and S5G).

**Figure 4.**
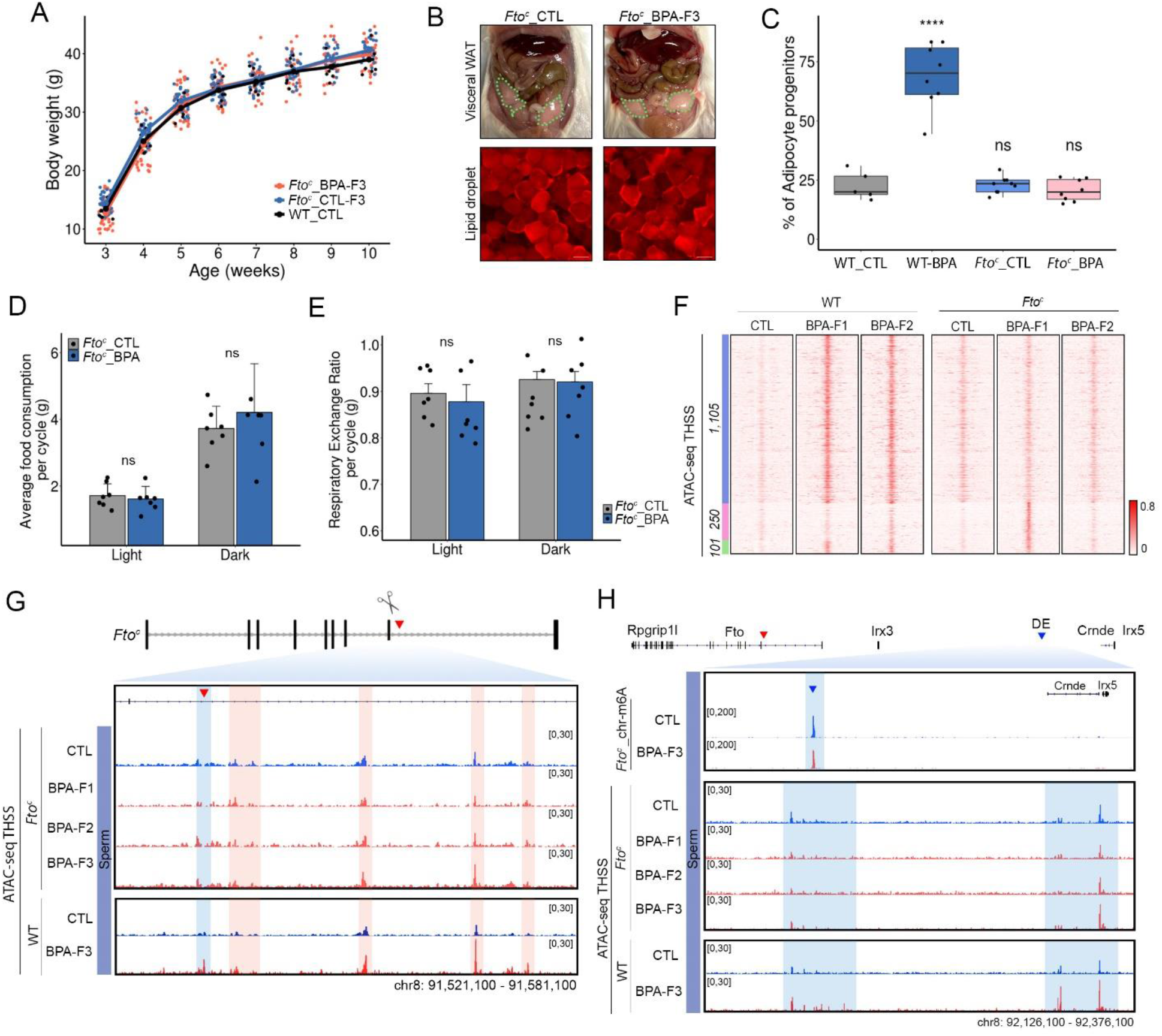
Deletion of the CTCF site in the *Fto* proximal enhancer prevents transgenerational transmission of obesity. (A) *Fto^c^* mice ancestrally exposed to BPA fail to gain weight with age with respect to unexposed *Fto^c^*_CTL and WT_CTL (*Fto^c^*_CTL n=34, *Fto^c^*_BPA n=40, WT_CTL n=9). (B) *Fto^c^* mice ancestrally exposed to BPA do not have excess visceral adipose tissue with respect to unexposed *Fto^c^*_CTL. Adipocytes accumulate the same amount of lipids and are the same size in unexposed and exposed *Fto^c^* mice. (C) *Fto^c^* mice ancestrally exposed to BPA have the same number of adipocyte progenitor cells in visceral adipose tissue as unexposed mice (*Fto^c^*_CTL n=9, *Fto^c^*_BPA n=8, WT_CTL n=5, WT_BPA n=8). (D) *Fto^c^* mice ancestrally exposed to BPA have the same food intake during the light and dark cycles as unexposed controls (*Fto^c^*_CTL n=7, *Fto^c^*_BPA n=7). (E) The respiratory exchange rate, which measures consumption of stored fats versus carbohydrates from a recent meal, is the same in *Fto^c^* mice ancestrally exposed to BPA and unexposed controls (*Fto^c^*_CTL n=7, *Fto^c^*_BPA n=7). (F) Comparison of chromatin accessibility determined by ATAC-seq in sperm of WT and *Fto^c^* mice unexposed and ancestrally exposed to BPA. Most sites induced by BPA exposure in WT mice do not increase in accessibility in *Fto^c^* mice (blue cluster). Only a small subset is induced in *Fto^c^* BPA-F1 mice, but they return to normal in the F2 generation (green cluster). Exposure to BPA results in increased accessibility at 250 sites (pink cluster) in *Fto^c^* but not WT mice, suggesting that deletion of the FOXA1/CTCF site in the proximal enhancer alters the response to BPA. These sites do not persist to the F2 generation. (G) The *Fto* proximal enhancer does not show increased chromatin accessibility and TF occupancy in the sperm of *Fto^c^* mice ancestrally exposed to BPA compared to controls. (H) Levels of m^6^A in the eRNA located in the *Fto* distal enhancer are not affected in *Fto^c^* mice ancestrally exposed to BPA. Chromatin accessibility and TF occupancy in the *Fto* distal enhancer is the same in the sperm of *Fto^c^* mice ancestrally exposed to BPA compared to controls. **** p<0.0001, ***p<0.001, ** p<0.01, *p<0.05, ns not significant.

The absence of a metabolic response to BPA exposure in mice carrying the *Fto^c^* deletion suggests a critical role for the *Fto* proximal enhancer in the transgenerational transmission of obesity. To gain additional insights into the mechanisms underlying this process, we performed ATAC-seq in sperm from *Fto^c^*_CTL and *Fto^c^*_BPA from the F1, F2, and F3 generations. Most sites induced by BPA exposure in the sperm of WT mice are not induced in sperm of *Fto^c^* males (Figure 4F, blue cluster), suggesting that these ATAC-seq peaks are a consequence of the activation of the *Fto* enhancer rather than BPA exposure. Only a small subset of sites induced in WT sperm are observed in F1 *Fto^c^* sperm, but these sites fail to be transmitted to the next generation and disappear in F2 sperm (Figure 4, green cluster). Interestingly, the deletion of the FOXA1/CTCF sites in *Fto^c^* mice results in the induction of 250 ATAC-seq peaks in the F1 generation that are not induced in WT animals and fail to persist in F2 (Figure 4, pink cluster). This observation suggest that suggests that lack of the FOXA1/CTCF sites results in an altered response to BPA, suggesting that non-coding variants affecting TF binding sites may make specific individuals differentially susceptible to environmental effects. The ATAC-seq peak containing the FOXA1 and CTCF motifs in the *Fto* proximal enhancer is absent in *Fto^c^* mice exposed to BPA (Figure 4G). Surprisingly, other ATAC-seq peaks in the *Fto* BPA proximal enhancer containing motifs for ESR1, AR, and PPARγ remain the same in *Fto^c^*_BPA as in *Fto^c^*_CTL and in WT_CTL, suggesting that the FOXA1/CTCF site is required for the full TF occupancy and presumed activation of this enhancer (Figure 4G). The FOXA1/CTCF site is also required for the activation of the distal enhancer (Figure 4H). Levels of m^6^A in chromosome-associated RNAs, including the eRNA present in the *Fto* BPA distal enhancer, do not change in F3 *Fto^c^*_BPA mice with respect to *Fto^c^*_CTL, and occupancy of CTCF and AR at sites detected by ATAC-seq remain the same as in control (Figure 4H).

### The *Lep* gene is activated in visceral fat tissue of mice ancestrally exposed to BPA

The obesity phenotype observed in mice ancestrally exposed to BPA is characterized by unregulated eating resulting in decreased utilization of fat as a fuel source. These changes could result from misregulation of expression of genes in the vicinity of the *Fto* locus due to activation of the proximal and distal enhancers in tissues of the adult responsible for the observed phenotypes. To test this possibility, we first performed ATAC-seq in liver tissue from control and exposed BPA-F4 male mice. The results show around 500 ATAC-seq peaks gained or lost between the two conditions (Figure 5A and Figure S6A) and these sites are located preferentially in intergenic regions and introns (Figure S6B). Genes adjacent to these sites encode proteins involved in metabolic processes known to be affected in the liver as a result of feeding and obesity (Figure S6C). Examples of these include *Cyp7a1*, which shows decreased accessibility and encodes a monooxygenase that catabolizes cholesterol to bile acids and is is normally upregulated during fasting; and *Fasn*, which shows increased accessibility and encodes an enzyme involved in the synthesis of fatty acids and it is normally upregulated after feeding (Figure 5B). Analysis of the region surrounding *Fto* indicates that the FOXA1/CTCF sites in the proximal enhancer in intron 8 are present in the liver tissue of control and BPA-F4 mice. Other ATAC-seq sites found in the proximal and distal enhancers are the same in the two conditions but they are different from those found in sperm (Figure S6A). These results suggest that the proximal FOXA1/CTCF sites become occupied in liver tissue during normal development, and that differential activation of these enhancers is not responsible for the observed chromatin changes in genes involved in lipid metabolism in the liver of BPA-F4 mice with respect to control. To further explore the origin of the observed phenotypes, we performed ATAC-seq in visceral adipose tissue from control, BPA-F4, overweight BPA-F6, and lean BPA-F6 males. The results show that the FOXA1/CTCF site in the *Fto* proximal enhancer is also present in both mature adipocytes and adipocyte progenitor cells, with no differences between control and obese mice (Figure S6A). Interestingly, the pattern of accessible sites indicates the binding of different TFs in different tissues to the same intronic enhancer in the *Fto* gene. In agreement with these observations, RNA-seq data indicates that the expression of *Fto* and adjacent genes is the same in fat tissue and adipocyte precursors from control and BPA-F4 animals (Figure S6D). These results suggest that the obesity observed in BPA-Fi animals is not due to deregulation of *Fto* in fat tissue or adipocyte progenitor cells.

**Figure 5.**
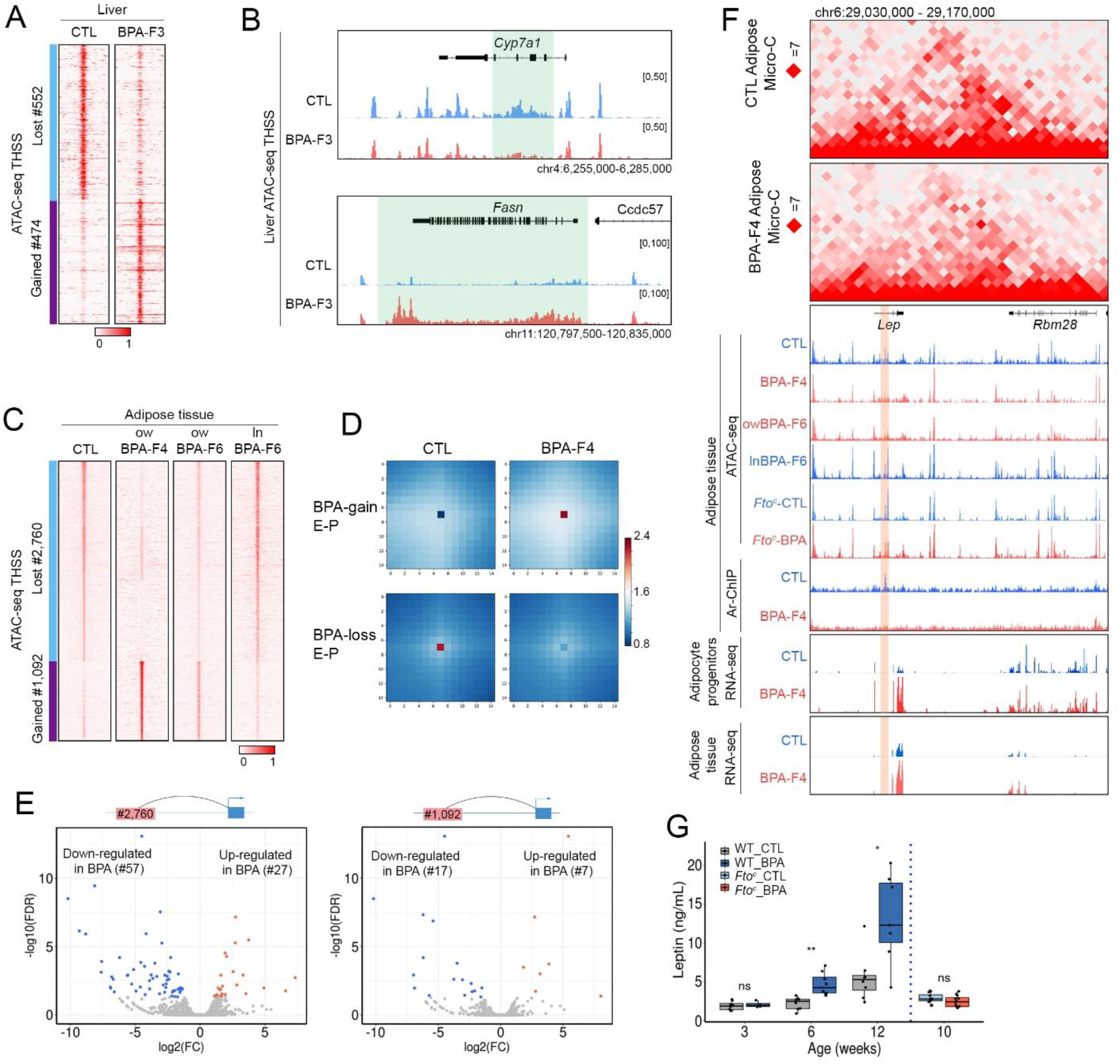
Changes in chromatin occupancy and 3D interactions in liver and visceral fat tissue. (A) Analysis of ATAC-seq results in liver of WT unexposed and BPA-F4 generation males. The cluster marked in blue contains sites lost in BPA-F4 mice whereas the cluster marked in purple contains sites gained. (B) Examples of specific genes where chromatin accessibility decreases (*Cyp7a1*, top) or increases (*Fasn*, bottom) in liver cells from WT BPA-F3 mice with respect to unexposed control. (C) Summary of differential chromatin accessibility results in visceral adipose tissue from control, BPA-F4, overweight BPA-F6, and lean BPA-F6. (D) Increased or decreased interaction frequency between enhancers (E) and promoters (P) located at differential THSSs based on Micro-C data from control and BPA-F4 visceral adipose tissue. (E) Up- and down-regulated interactions between differential THSSs and their target promoters based on Micro-C data from control and BPA-F4 visceral adipose tissue. (F) Micro-C heatmaps, ATAC-seq, RNA-seq, and AR ChIP-seq data in visceral adipose tissue of control, BPA-F4, BPA-F6 and *Fto^c^* mice in the region surrounding the *Lep* gene. The lower panels show RNA-seq in the same region from adipose tissue and adipocyte progenitor cells. (G) Levels of leptin at different ages in serum from WT and *Fto^c^* mice unexposed or ancestrally exposed to BPA (3 weeks; WT_CTL n=8, WT_BPA n=4, 6 weeks; WT_CTL n=8, WT_BPA n=8, 12 weeks; WT_CTL n=8, WT_BPA n=7, 10 weeks; *Fto^c^*_CTL n=12, *Fto^c^*_BPA n=12). **** p<0.0001, ***p<0.001, ** p<0.01, *p<0.05, ns not significant.

It is possible that miss-expression of *Fto* earlier in development of BPA-Fi mice may have altered expression of other genes that in turn are responsible for changes in fat tissue differentiation and gene expression, which cause the obesity phenotype in the adult. We thus compared ATAC-seq data obtained in visceral adipose tissue from control, BPA-F4, and lean and over-weight BPA-F6 male mice. We find dramatic changes in the distribution of accessible chromatin sites, with 1,092 sites increased or gained and 2,760 decreased or lost (2-fold change, p value ≤0.05) in the adipose tissue of BPA-F4 and overweight BPA-F6 with respect to control and lean BPA-F6 (Figure 5C). TF sites whose binding is altered in adipose tissue are located almost exclusively in intergenic and intronic regions of the genome, suggesting that they may correspond to enhancers (Figure S6E), and contain motifs for a variety of TFs, including several nuclear hormone receptors. None of the altered TF binding sites found in adipose tissue correspond to altered TF sites observed in sperm. To further examine the basis for the obese phenotype, we performed Micro-C using adipose tissue of control and BPA-F4 males (see Table S1 for quality control of Micro-C reads). We then identified significant interactions using Fit-Hi-C and identified those involving TF sites from ATAC-seq data altered in BPA-F4 versus control. TF sites gained in BPA-F4 interact with increased frequency with target genes whereas those lost or decreased interact with lower frequency (Figure 5D). We then examined RNA-seq data obtained in adipose tissue of BPA-F4 and control males and identified target genes of altered TF sites that increase or decrease in BPA-F4 adipose tissue with respect to control (Figure 5E). We also used RNA-seq to identify genes whose expression is altered in adipocyte progenitors obtained from fat tissue of BPA-F4 versus control. The expression of *Fto* or surrounding genes is not altered in fat tissue of obese animals with respect to unexposed controls. One of the genes up-regulated in both adipocyte precursors and visceral fat tissue of BPA-F4 is *Lep*, (Figure 5F, bottom panels) which encodes the leptin hormone involved in the regulation of body weight (*39*). Increased levels of *Lep* RNA correlate with increased levels of leptin protein in the blood of BPA-F4 mice. Levels of leptin are initially the same in control and BPA-F4 at 3 weeks, but leptin levels increase dramatically with age in BPA-F4 males (Figure 5G). Upregulation of *Lep* RNA in BPA-F4 animals correlates with the disappearance of an accessible region in the intron of the *Lep* gene. This site, which contains the motif for AR, is present in fat tissue of control and lean BPA-F6 males but absent in BPA-F4 and overweight BPA-F6 animals (Figure 5F). ChIP-seq with antibodies to AR confirms the presence of this protein at this site in adipose tissue of control and its absence in obese males (Figure 5F). The loss of AR from this putative silencing element correlates with a decrease of interactions between *Lep* and the adjacent *Rbm28* gene (Figure 5F). Contrary to WT mice, the ATAC-seq site corresponding to the presence of AR does not decrease in *Fto^c^* mice exposed to BPA (Figure 5F), and these mice do not have higher levels of leptin protein in blood compared to controls of the same age (Figure 5G). These observations indicate that mice ancestrally exposed to BPA produce higher levels of leptin protein in visceral fat tissue via increased expression of the *Lep* gene, and this requires the FOXA1/CTCF sites in the proximal enhancer of the *Fto* gene. However, CTCF occupancy of this enhancer is the same in the liver, adipocyte progenitors, and fat tissue of control and BPA-F4 mice, suggesting the effect of differential occupancy observed in sperm takes place earlier in development or it acts in a different tissue, which is primarily responsible for the obese phenotype with secondary indirect effects on other target organs.

### BPA exposure affects the differentiation of AgRP/NPY neurons in the hypothalamus

Mice ancestrally exposed to BPA display increased food intake suggestive of enhanced appetite, which is controlled by anorexigenic POMC and orexigenic AgRP neurons located in the arcuate nucleus (ARC) of the hypothalamus, which integrate signals from the satiety hormone leptin (*40*). BPA mice have increased levels of leptin but behave as if they have become resistant to this hormone. POMC and AgRP neurons are formed postnatally from tanycytes and radial glia-like neural stem cells (RG-NSCs), and their differentiation is regulated by *Irx3* and *Irx5* (*41*). As described above, these two genes are contacted by the *Fto* proximal and distal enhancers at higher frequency in sperm of mice ancestrally exposed to BPA. To examine whether BPA exposure affects the formation of POMC or AgRP neurons, we microdissected the region around the hypothalamic arcuate-median eminence (ARC-ME) (Figure S7A) and performed single-nucleus ATAC-seq (snATAC-seq) for two biological replicates of WT unexposed and BPA-F3 adult mice. Using the ArchR software suite (*42*), we obtained 19,059 single-nucleus epigenomes after quality control filtration. Harmony was used for batch effect correction (*43*), iterative latent semantic indexing was performed for dimensionality reduction, and clusters were identified with Seurat (*44*) (Figure S7B-C, Table S2). Using known hypothalamic cell-type specific markers (*41, 45–47*), we first identified 4 clusters containing oligodendrocytes and oligodendrocyte progenitor cells (NG2-OPCs), ME non-neuronal cells, and neurons (*Tubb3*^+^) (n=19,059 nuclei) (Figure S7C-D). Neuronal cells were then clustered into 25 distinct clusters (Figure S7E) and further sub-clustered after eliminating cells expressing markers of ventromedial hypothalamus (VMH) (*Sf1^+^*, *Nr5a1^+^, Fezf1^+^*) (Figure S7E-F). Clusters were then assigned to known ARC-ME neurons based on gene activity scores compiled from transposase-accessibility signals in the vicinity of key cell type-defining genes (Figure 6A, S7F and S8, Methods). The cell types identified in the unsupervised analysis using snATAC-seq are consistent with previous studies profiling single-cell transcriptomes of the ARC-ME region of the hypothalamus (*45*).

**Figure 6.**
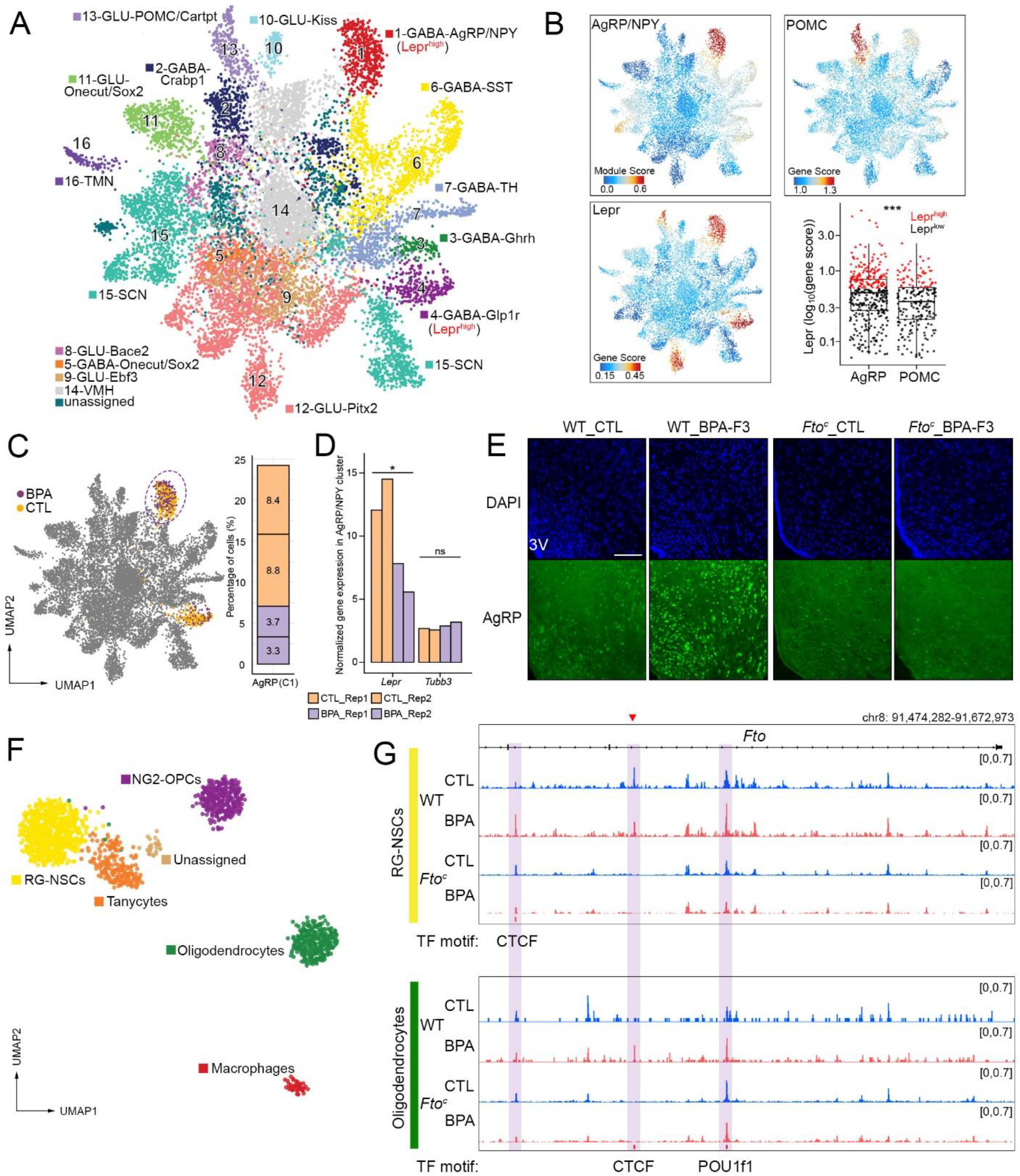
Mice ancestrally exposed to BPA have abnormal ratios of AgRP/NPY neurons in the hypothalamus. (A) Uniform manifold approximation and projection (UMAP) representation of snATAC-seq data (n = 8880 CTL nuclei, n = 4348 BPA-F3 nuclei) colored by neuronal clusters cell types. ARC-ME neurons (clusters 1-13) and neurons of the suprachiasmatic nucleus (SCN), tuberomammillary nucleus (TMN), and ventromedial hypothalamus (VMH) are shown (clusters 14-16). (B) Gene scores for *Lepr* and *Pomc*, and gene module score for AgRP/NPY (gene module: *Npy*, *Agrp*, *Slc32a1*) rendered with imputation (*54*). Also shown is a box plot of the *Lepr* gene score in AgRP/NPY and *Pomc*/*Cartpt* nuclei from clusters 1 and 13. ((C) Left: UMAP plot of CTL and BPA-F3 nuclei with high *Lepr* gene score (Clusters 1 and 4). Right: Percentage of orexigenic (*Agrp*^+^/*Npy*^+^, cluster 1, *LepR^high^*) neurons out of total non-VMH neurons in each biological replicate of CTL and BPA. (D) Pseudo-bulk normalized expression of LepR RNA in snRNA-seq data in AgRP/NPY neurons (clusters 1) in each biological replicate of CTL and BPA (DEseq2 normalized counts x10^3^ are shown). (E) Sections of brain regions proximal to the arcuate nucleus of the hypothalamus from WT and *Fto^c^* mice unexposed or ancestrally exposed to BPA and stained with DAPI and AgRP peptide antibodies. (F) Subclustering of non-neural lineages colored by major cell types found in hypothalamic ARC-ME proximal regions. (G) Differential chromatin accessibility at the *Fto* region surrounding introns 7 and 8. Pseudo-bulk accessibility in snATAC-seq datasets of RG-NSCs and oligodendrocyte clusters in exposed and unexposed WT and *Fto^c^* mice (FDR<0.05, log_2_(FC)>1); differential peaks are highlighted.

Cluster 1 neurons have high gene activity scores for both *AgRP* and *NPY* whereas neurons in cluster 13 show high gene activity scores for *Pomc* and *Cartpt* (Figure 6B). Both clusters have high gene activity scores for the leptin receptor (*Lepr*) gene. However, the gene activity score for *Lepr* is lower in POMC neurons and, as has been previously shown, only around 12% of POMC neurons express LepR (*48*), making AgRP/NPY neurons more likely candidates to transduce the effect of the high Lep levels observed in BPA mice (Figure 6B). We then compared snATAC-seq data from WT BPA-exposed and unexposed animals and found that BPA-F3 mice have more AgRP/NPY neurons than unexposed animals (Figure 6C). To explore how this excess of AgRP/NPY neurons may transduce the effects of high leptin levels found in BPA mice, we examined *Lepr* RNA expression from the same nuclei of pseudo-bulk AgRP/NPY neurons in snRNA-seq data of WT BPA-exposed and unexposed animals. We found that BPA-F3 mice express significantly less LepR RNA (Figure 6D; FDR<0.05, Fold change > 2). Given the high level of leptin found in BPA mice, the reduction of LepR in AgRP/NPY neuron provides an explanation for how these mice are resistant to high circulating leptin levels, similar to mice with a deletion of LepR in AgRP neurons or homozygous for the diabetes spontaneous mutation *Lep^db/db^* (*49, 50*). To confirm that BPA-F3 mice have more AgRP neurons than unexposed controls, we performed immunofluorescence microcopy in the ARC region of the hypothalamus in separate mice using antibodies against the AgRP peptide. Results show an increase in the number of AgRP neurons in BPA-F3 compared to WT unexposed mice, and these neurons have higher levels of AgRP (Figure 6E). Importantly, the number of AgRP neurons does not increase in *Fto^c^* mice carrying the deletion of the FOXA1/CTCF site (Figure 6E), supporting the conclusion that AgRP neurons may be responsible for the increased food intake and obesity phenotype in mice ancestrally exposed to BPA.

AgRP neurons arise postnatally from non-neuronal cells in the ARC-ME, including tanycytes and RG-NSCs which, together with oligodendrocytes, have also been shown to play important roles in the regulation of leptin responses, food intake and metabolisms (*41, 46*). Differentiation of AgRP neurons is controlled by *Irx3* and *Irx5*, (*41*) whose expression is regulated by enhancers in the *Fto* gene (*51, 52*). However, no changes in chromatin accessibility are observed at the *Fto* proximal or distal enhancers in AgRP neurons (Figure S9A). Since these neurons differentiate from non-neuronal cells in ARC-ME, we examined snATAC-seq information for these non-neuronal cell types. We identified five non-neuronal cell types in unsupervised analyses (Figure 6F, S9B, and Table S2) and examined possible changes in chromatin accessibility between BPA-exposed and non-exposed mice. RG-NSCs (*Agt^+^*, *Ntsr2 ^+^*, *Hest5*^+^ and low gene score for *Gfap*) do not show changes in ATAC-seq signal at the proximal *Fto* enhancer. However, a new CTCF site located in intron 7 becomes more accessible in WT BPA mice compared to unexposed controls (FDR<0.05, FC>2) (Figure 6G). To examine whether this new CTCF site in intron 7 is associated with the obesity phenotype, we performed snATAC-seq using the ARC-ME and proximal hypothalamic regions from *Fto^c^* mice and the data was processed using the same methodology described above to identify non-neuronal cells present in the ARC-ME of *Fto^c^* mice (Figure S10 and Table S2). We find that the CTCF site induced in WT RG-NSCs by ancestral exposure to BPA does not increase in accessibility in *Fto^c^* mice (Figure 6G). In addition to RG-NSCs, mature oligodendrocytes (oligodendrocytes; *Mag^+^, Opalin^+^, Mal^+^*) exhibit an increase in accessibility (FDR<0.05; FC>2) at the CTCF site located in the proximal *Fto* enhancer as well as at a site containing the binding motif for POU1f1 (Figure 6G). These changes do not take place in *Fto^c^* mice (Figure 6G), suggesting that they may be involved in eliciting the obesity epiphenotype. However, the increased accessibility of the *Fto* proximal enhancer in oligodendrocytes does not correlate with changes in chromatin accessibility or gene activity scores at *Irx3* and *Irx5*, in the adult brain. These changes may take place postnatally when the arcuate nucleus matures and neurons involved in appetite control first differentiate (Figure S9C) (*41*).

## Discussion

Evidence for adverse health effects in humans resulting from ancestral exposures to environmental changes is extensive. For example, studies of the Överkalix cohorts exposed to food shortages in northern Sweden have shown that paternal grandfather’s food access in pre-puberty predicts all-cause mortality in male grandchildren (*53*). An overwhelming number of similar reports in humans and laboratory animals suggests that the mammalian germline is exquisitely sensitive to environmental effects that can be passed onto future generations. Although some of these effects, such as diet or stress, have been encountered by humans for thousands of years and can thus be argued to be “natural”, and may have played a role in adaptation during evolution, changes in the chemical composition of the environment since the industrial revolution have resulted in the exposure of the human germline to a range of chemicals not experienced before. Some of these chemicals have structures similar to sex hormones and they may exploit existing signaling pathways with normal roles in germline differentiation to produce changes in the germ cell epigenome that result in adverse health effects in subsequent generations. In spite of the critical importance of this problem and the wealth of available information, there is currently a lack of mechanistic molecular explanations for how environmentally-induced alterations in the mammalian germline epigenome can be transmitted through multiple generations. Observations reported here suggest that exposure to BPA results in alteration of the male and female germline epigenomes manifested by new binding sites for a variety of transcription factors. Many of these sites correspond to nuclear hormone receptors and this may be due to the ability of BPA to bind to these proteins. Thus, it is likely that exposure to different types of environmental perturbations will elicit changes in other classes of transcription factors, resulting in the misregulation of distinct groups of genes and giving rise to different phenotypes. If this is the case, it may be possible to assign specific exposures to distinct TF distribution signatures and to understand the additive or synergistic effects of complex exposures. This may allow the establishment of rules that can translate TF signatures to complex exposures, making adverse health effect predictable.

*Fto* enhancers activated by ancestral BPA exposure contain binding sites for various nuclear hormone receptors. It is possible that BPA binds to these receptors in PGCs, which then can bind to chromatin at the time of the exposure when DNA is demethylated. The presence of these factors bound to chromatin may then interfere with DNA remethylation during germline development after E13.5. We have previously shown that a subset of TFs bound to the sperm genome are also found on chromatin during preimplantation development (*12*). Therefore, new patterns of TF occupancy in the gametes caused by environmental exposures could persist in the embryo after fertilization, where they may again protect their binding sites from remethylation after the blastocyst stage. Once their binding sites are unmethylated, TFs may not need to be bound to DNA continuously in order to maintain a memory state of accessible chromatin. Although such a mechanism may explain transgenerational inheritance in general, the involvement of *Fto* in the case of BPA exposure and the role of the FTO protein in m^6^A methylation offers an additional or alternative explanation for the transgenerational transmission of obesity. It is possible that activation of the *Fto* intronic enhancer by BPA results in lower levels of m^6^A and preferential persistence of the chromatin-associated eRNA from the intergenic enhancer after fertilization. Since this distal intergenic enhancer interacts with the *Fto* promoter and its intronic enhancer, this could lead to activation of *Fto* during preimplantation development. The positive feedback loop between the two enhancers could contribute to the continued activation of the proximal and distal enhancers until the epiblast stage when the germline differentiates and the transmission of this active state between generations.

Results presented here suggest that the FOXA1/CTCF site present in the *Fto* proximal enhancer is required to elicit the BPA-dependent obesity epiphenotype and its transgenerational transmission. The role of this CTCF site may be to target the adjacent enhancers to the appropriate promoters during development. The FOXA1/CTCF site appears to be occupied in several adult tissues, including liver, adipocyte progenitors, and mature adipocytes, of unexposed mice, suggesting that it plays a role during normal development and that these tissues may not be the primary target of missactivation of *Fto* enhancers in mice ancestrally exposed to BPA. Since these mice appear to become obese due to unregulated eating, the primary target of BPA-dependent *Fto* enhancers may be the ARC region of the hypothalamus involved in appetite control. In support of this hypothesis we find an increased number of orexigenic AgRP neurons with lower levels of *LepR* RNA that could signal BPA mice to increase their food intake in spite of the high circulating Lep levels. However, these cells do not show differences in chromatin accessibility in the *Fto*, *Irx3*, and *Irx5* genes. Instead, these differences are found in their RG-NSC precursor cells and in mature oligodendrocytes. The involvement of these cells in the control of appetite has not been studied extensively but it has been shown previously that the formation of mature oligodendrocytes located in the ME region can be regulated in response to nutritional signals, and the mature oligodendrocytes located in the ME region can in turn control the access of circulating metabolic hormones, including leptins, to the rest of the ARC-ME (*46*).

Together, these observations suggest that environmentally-induced alterations in the gametes can be transmitted to the embryo after fertilization and can be maintained in a subset of somatic cells of the adult to elicit specific phenotypes. These findings offer a molecular mechanistic interpretation of numerous studies showing adverse health outcomes after different types of environmental exposures in humans and laboratory animals. The finding that both genetic mutations and environmentally induced epigenetic alterations in the *Fto* gene lead to similar phenotypic outcomes suggests that much of the missing genetic variation required to explain many human diseases may be epigenetic in origin.

## Supplemental Materials

### Methods

#### Experimental model and BPA administration

Mice were maintained and handled in accordance with the Institutional Animal Care and Use policies at Emory University. All experiments were conducted according to the animal research guidelines from NIH and all protocols for animal usage were reviewed and approved by the Institutional Animal Care and Use Committee (IACUC). Mice were housed in standard cages on a 12: 12 h light:dark cycle and given ad lib access to food and water. Healthy 8-week-old CD1 mice (Charles River Labs) not involved in previous procedures were used for all experiments. Gestating females (F0) were administered daily intraperitoneal injections of Bisphenol A (Sigma 239658, 50 mg/kg) or sesame oil (Sigma S3547). Injections were performed from embryonic day 7.5 through 13.5. No sibling breeding was used to avoid inbreeding artifacts. To examine transgenerational transmission of obesity, two pregnant females were exposed to BPA and two additional ones were injected with sesame oil vehicle as controls. Crosses between the F1 progeny of each female were performed as shown in Figure S1A. This experiment was repeated 7 times by three different investigators. Only two of the experiments were carried out to reach the F7 generation. Numbers of animals used from each generation to perform experiments described in the Results section are described in the corresponding figure legends. The body weight data shown in the figures is a combination of that obtained in all 7 experiments performed. The number of animals decreases with generation or with age because some animals were sacrificed to perform experiments described in the manuscript. Although the weight of all animals in all experiments was measured, only a subset whose weight was determined at specific time points is shown in the figures.

#### Isolation of mouse sperm and oocytes

Euthanasia was performed by cervical dislocation and the epididymis was removed. Mature sperm were collected from the dissected cauda epididymis of 8-10-week-old CD1 mice (Charles River Labs). After dissection to eliminate blood vessels and fat, the cauda epididymis was rinsed with PBS, deposited in Donners medium in a cell culture plate, and punctured with a needle. Sperm were then transferred to a tube and allowed to swim up for 1 hr (*1*). Purity of sperm was determined by examination under a microscope after DAPI staining. After counting 1000 sperm, purity was determined to be at least 99.9% if no contaminating cells were observed. Ovaries were harvested from 16-day-old CD1 female pups for the preparation of GV-stage oocytes. The zona pellucida was removed with hyaluronidase to avoid any residual cumulus cells.

#### Isolation of adipocyte progenitor cells

Visceral adipose tissue (1g) was minced manually in PBS (Ca2+ and Mg2+ free), transferred to 10 ml of DMEM/F12 containing 0.8% fatty acid–free bovine serum albumin, collagenase D (1.5 units/ml; Roche), and dispase II (2.4 units/ml; Roche), and then incubated at 37°C with agitation for 45 min. The reaction was quenched with DMEM/F12 containing 10% FBS and dissociated cells were passed through a 100 μm filter. After centrifugation, the pellet was resuspended in red blood lysis buffer and incubated for 5 min at RT. The reaction was quenched with DMEM/F12 containing 10% FBS and passed through a 40 μm filter. After recovering cells in chilled MACS buffer, adipocyte progenitor cells were isolated using the Adipose Tissue Progenitor Isolation Kit, mouse (Miltenyi Biotec, 130-106-639) according to the manufacturer’s protocol.

#### *In vitro* fertilization (IVF) and embryo transfer

IVF was performed using BPA-F2 MII oocytes and epididymal sperm from control males. In brief, superovulation was induced in BPA-F2 CD1 females at 6-8 weeks of age using intraperitoneal injection of 7.5 IU of pregnant mare serum gonadotropin (PMSG) followed by 5 IU of human chorionic gonadotropin (hCG) at 48 h post-PMSG injection. MII oocytes were harvested at 13-14 h post-hCG injection. Sperm collected from the cauda epididymis were transferred into droplets containing oocytes. After 3-4 h of co-incubation, oocytes were freed from sperm and cumulus cells using a fine glass pipette. For embryo transfer experiments, two-cell embryos produced by IVF were transferred into the oviducts of 7 pseudopregnant control CD1 females mated with a vasectomized male. A total of 42 pups were obtained and analyzed as described in the Results.

#### Targeted deletion of a FOXA1/CTCF binding site in the *Fto* proximal promoter

To generate mice carrying a deletion of the FOXA1 and CTCF binding sites in intron 8 of the *Fto* gene, two clustered regularly interspaced short palindromic repeat (CRISPR) guide RNAs (gRNAs) were designed as shown in Figure 5SA. 50 ng/μl of each gRNA and 100 ng/μl of Cas9 RNA were injected into one-cell CD1 zygotes. Embryos were cultured to the 2-cell stage overnight and transferred to pseudopregnant females. After birth, all mice were genotyped by PCR amplification using genomic DNA isolated from mouse tails with PCR primers (5’-CATCAAAGTTGAGCCTCCAAG -3’ and 5’-CATCTCAGGAGCCAACAAAGG-3’) followed by sequencing of the amplified DNA to verify the targeted deletion. Homozygous mutant mice were isolated using appropriate crosses.

#### Indirect calorimetry and food intake

Control- and BPA-F4 CD1 mice (n=7-13 per group) were singly housed in metabolic cages (Columbus Instruments CLAMS-HC) and maintained under otherwise standard housing conditions (12-hour light-dark cycle; 22.2 ± 1.1°C) for indirect calorimetry and food intake assessments. After a 2-day acclimation period, O_2_ consumption, CO_2_ production, and food intake were monitored over 14-minute periods for consecutive light-dark cycles over 3 successive days. The respiratory exchange ratio (RER) was calculated as the ratio of CO_2_ production over O_2_ consumption in 14-minute intervals. Mice were provided ad libitum food access, and food intake was averaged for each mouse during each 12-hour cycle (light or dark).

#### Determination of BPA levels by mass spectrometry

F0 pregnant females were injected with 50 mg/kg of BPA or sesame oil vehicle every 24 h from E7.5 to E13.5 of fetal development. Embryos from pregnant females were collected 10 min, 2 h, and 24 h post injection on E13.5. From each timepoint, 1 g of embryo tissue was pooled from 5 embryos, homogenized in 500 μl of water, and BPA was extracted as described (*2*). Briefly, 0.25 mg of BPA-d16 was spiked in as an internal standard, BPA was extracted with 500 μl of acetonitrile, quenched with 100 mg of MIX I and 100 mg of MIX IV (*2*). The upper organic layer was then collected and evaporated under a stream of nitrogen gas. The precipitate was dissolved in 200 μl of water/methanol (70/30 v/v). Negative ion LTQ-FTMS was run with isocratic 50% acetonitrile and 50% 20 mM ammonium formate/water solution, with a retention time of 3.1 minutes and 5 μg/ml limit of detection.

#### Measurement of serum leptin

Mice were exsanguinated at time of death by cardiac puncture to collect whole blood. Blood was allowed to clot in microfuge tubes at room temperature for 30 minutes, and then centrifuged at 1,500 g for 15 minutes at 4°C to collect serum. Serum leptin levels were assayed using a Millipore mouse leptin ELISA kit (Cat. # EZML-82K) according to the manufacturer’s instructions. The sensitivity of the ELISA assay was determined by the manufacturer to be 0.05 ng/ml for a 10-μl sample size, and the intra-assay coefficient of variation was determined to be between 1.06 and 4.59 %.

#### Lipid droplet fluorescence assay

Visceral adipose tissue was fixed for 30 min in 4% paraformaldehyde, washed with PBS, and stained with HCS LipidTox detection reagent (Invitrogen H34476). Stained tissues were then photographed under a Zeiss Axio Observer Z1 microscope.

#### Assay for transposase-accessible chromatin using sequencing (ATAC-seq)

ATAC-seq was carried out using the Omni-ATAC protocol (*3*). After cells were counted, the nuclei from 100,000 sperm or 50,000 adipocyte progenitor cells were isolated with Lysis Buffer (10 mM Tris-HCl pH 7.4, 10 mM NaCl, 3 mM MgCl_2_) containing 0.1% NP40, 0.1% Tween-20, and 0.01% digitonin. The purified nuclei pellet was then resuspended in the transposase reaction mix containing 0.05% digitonin and incubated for 30 min at 37°C. To perform ATAC-seq using GV oocytes, around 500 oocytes were deposited in a tube with 500 μl of RSB (10 mM Tris-HCl, 10 mM NaCl and 3 mM MgCl_2_), and spun down at 500 rcf for 5 min. After careful removal of the supernatant, 15 μl of 2x TD buffer, 0.75 μl of Tn5, 0.3% of 1% Digitonin, 0.3% of 10% Tween20 and 0.3 μl of 10% NP40 were added, and samples were incubated for 30 min at 37°C. Following incubation, nuclei were treated with Proteinase K at 55°C for 2 h, and gDNA was isolated by phenol:chloroform:isoamyl alcohol and EtOH precipitation. ATAC-seq with liver and fat tissue were performed using the ATAC-seq protocol for frozen tissues (3). Tissues were prepared in a 2 ml Dounce homogenizer containing 500 μl of homogenization buffer. Nuclei were then collected by Iodixanol gradient and ATAC-seq reactions with 50,000 nuclei were performed as described above.Library amplification was done with 2x KAPA HiFi mix (Kapa Biosystems) and 1.25 µM indexed primers using the following PCR conditions: 72°C for 5 min; 98°C for 30 s; and 10-11 cycles at 98°C for 10 s, 63°C for 30 s, and 72°C for 1 min. Libraries were paired-end sequenced on Illumina Hiseq2500 v4 or NovaSeq 6000 instruments.

#### Bisulfite Sanger sequencing

DNA was extracted from epididymal sperm using phenol/chloroform/isoamylalcohol (25:24:1). About 2 μg of DNA was bisulfite converted and purified using the Zymo EZ DNA methylation lightning kit (D5030) following the manufacturer’s protocol. Bisulfite converted DNA was then used as template for PCR amplification with KAPA HiFi HotStart Uracil kit (Roche, KK2801) using primers surrounding Site 1 in intron 8 of the *Fto* gene. The resulting 470 bp fragment was purified using the Zymoclean Gel DNA Recovery kit (D4008). Primer sequences used were as follows: forward 5’- TGGAGTAGGYGTTTTGAGGTGAAAGGGTAG-3’ and reverse, 5’- TCACTACRATTTTTCCTAACATAACAAAC-3’. Amplicons were ligated to T-Vector pMD19 (Takara, No.3271) with T4 ligase (NEB, M0202L) and transformed into DH-5a competent cells. DNA was isolated from monoclonal bacterial colonies and sequenced.

#### Whole genome bisulfite sequencing

DNA (1∼2 μg) was isolated from cauda epididymis sperm using phenol:chloroform:isoamyl alcohol followed by EtOH precipitation and spiked with 0.5% unmethylated lambda genomic DNA (Promega) to determine conversion efficiency after bisulfite treatment. The DNA was sheared with a Diagenode Bioruptor to yield DNA fragments ranging from 200 to 500 bp. After purification of the fragmented DNA using phenol/chloroform, DNA fragments were end repaired, A-tailed, and ligated to methylated Illumina adaptors. The Zymo EZ DNA methylation lightning kit (D5030) was used for bisulfite conversion and subsequent purification according to the manufacturer’s instructions. Two independent biological replicates per sample were then sequenced using paired-end 50 bp on Illumina Hiseq2500 v4 or NovaSeq 6000 instruments.

#### ChIP-seq

Sperm or somatic cells were crosslinked with 1% formaldehyde in 1xPBS for 10 min at RT, and the reaction was quenched with 125 mM glycine for 10 min at RT. After washing with PBS, cells were lysed with 5 mM PIPES, 85 mM KCl, 0.5% NP40 and 1x protease inhibitors (P8107S, NEB) on ice for 15 min. After centrifugation, cells were resuspended in RIPA buffer (1x PBS; 1% NP40; 0.5% sodium deoxycholate; 0.1% SDS; 1x protease inhibitors) and incubated on ice for 20 min. The purified chromatin was sonicated to 300-500 bp using a Diagenode Bioruptor. After 25 cycles (30 seconds on and 60 seconds off), the supernatant containing sheared chromatin was collected. Immunoprecipitation was performed overnight at 4°C with antibodies to the androgen receptor (sc-816, Santa Cruz) or CTCF (#3418, Cell Signaling Technology). Libraries for Illumina sequencing were constructed using the following standard protocol. Fragment ends were repaired using the NEBNext End Repair Module and adenosine was added at the 3’ ends using Klenow fragment (3’ to 5’ exo minus, New England Biolabs). Precipitated DNA was incubated with adaptors at room temperature for 1 hr with T4 DNA ligase (New England Biolabs) and amplified with Illumina primers. Two independent biological replicates per sample were then sequenced using paired-end 50 bp on Illumina Hiseq2500 v4 or NovaSeq 6000 instruments.

#### RNA-seq

Total RNA was isolated from epididymal sperm, visceral fat tissue, or adipocyte progenitor cells using Trizol reagent (Invitrogen) and ribosomal RNA was removed using the RiboMinus Transcrptome isolation kit (Invitogen, K1550). RNA concentration was measured using the Qubit RNA HS Assay kit (Thermo Fisher) and fragmented randomly by adding fragmentation buffer. cDNA was synthesized using RNA template and random hexamer primers. After terminal repair, A ligation, and sequencing adaptor ligation, the double-stranded cDNA library was completed by size selection and PCR enrichment. Illumina libraries were sequenced paired-end 50 bp on Illumina Hiseq2500 v4 or NovaSeq 6000 instruments. Two independent biological replicates per sample were then sequenced using paired-end 50 bp on Illumina Hiseq2500 v4 or NovaSeq 6000 instruments.

#### MeRIP-seq

Total RNA from sperm was isolated as described above. For m^6^A immunoprecipitation, purified RNA was digested using RNA Fragmentation Reagents (Ambion AM8740) by incubation at 70°C for 5 min in fragmentation buffer and precipitated via standard ethanol precipitation. Anti-m^6^A antibody (Synaptic Systems Cat.no 202 003) was incubated with 30 μl of Dynabeads Protein A for 1h at 4°C. Fragmented RNA was incubated with antibody-bead mixture overnight at 4°C. Library preparation was performed using the SMARTer Stranded Total RNA-seq kit v2 (Takara) according to the manufacturer’s protocol. Two independent biological replicates per sample were then sequenced using paired-end 50 bp on Illumina Hiseq2500 v4 or NovaSeq 6000 instruments.

#### Chromosome-associated RNA m^6^A-seq

Fractionation of chromosome-associated RNAs and generation of m^6^A-seq libraries were carried out as described (*4*). Briefly, 10^8^ sperm cells were isolated from the cauda epididymis and resuspended in cold lysis buffer (10 mM Tris-HCl, 0.05% NP40, 150 mM NaCl). After incubation on ice for 5 min, cells were resuspended in 2.5 volumes of sucrose solution (24% sucrose in lysis buffer), and then centrifuged at 4°C at 14,000 g for 10 min. The supernatant containing the cytoplasmic fraction was removed and the nuclei pellet was resuspended in glycerol buffer (20 mM Tris-HCl, 75 mM NaCl, 0.5 mM EDTA, 0.85 mM DTT, 0.125 mM PMSF, 50% glycerol) and an equal volume of cold nuclei lysis buffer (10 mM HEPES, 10 mM DTT, 7.5 mM MgCl_2_, 0.2 mM EDTA, 0.3 M NaCl, 1M urea, 1% NP40). After resuspending the pellet by vortexing, the nuclei were incubated on ice for 2 min and centrifuged at 4°C at 14,000 g for 2 min. The supernatant containing the soluble nuclear fraction/nucleoplasm was removed and the chromosome-associated fraction was collected in the pellet. Total RNA from the chromosome-associated fraction was isolated with Trizol. Ribosomal RNA was removed using the RiboMinus transcriptome isolation kit (Invitrogen, K1550), and then RNA was digested using RNA Fragmentation Reagents (Ambion AM8740) by incubation at 70°C for 5 min. m^6^A-immunoprecipitaion was performed using the EpiMark N6-Methyladenosine Enrichment Kit (NEB E1610S), followed by library preparation using the SMARTer Stranded Total RNA-seq kit v2 (Takara) according to the manufacturer’s protocols. Two independent biological replicates per sample were then sequenced using paired-end 50 bp on Illumina Hiseq2500 v4 or NovaSeq 6000 instruments.

#### In-situ Hi-C

In-situ Hi-C libraries were prepared using DpnII restriction enzyme as previously described (*5*). Briefly, 10 million sperm were crosslinked with 1% formaldehyde, quenched with glycine, washed with PBS, and permeabilized to obtain intact nuclei. Nuclear DNA was then digested with DpnII, the 5’-overhangs were filled with biotinylated dCTPs and dA/dT/dGTPs to make blunt-end fragments, which were then ligated, reverse-crosslinked, and purified by standard DNA ethanol precipitation. Purified DNA was sonicated to 200-500 bp small fragments and captured with streptavidin beads. Standard Illumina TruSeq library preparation steps, including end-repairing, A-tailing, and ligation with universal adaptors were performed on beads, washing twice in Tween Washing Buffer (5 mM Tris-HCl pH 7.5, 0.5 mM EDTA, 1M NaCl, 0.05% Tween 20) between each step. DNA on the beads was PCR amplified with barcoded primers using KAPA SYBR FAST qPCR Master Mix (Kapa Biosystems) for 5∼12 PCR cycles to obtain enough DNA for sequencing. Libraries were paired-end sequenced on an Illumina NovaSeq 6000 instrument. Two biological replicates were generated, and replicates were combined for all analyses after ensuring high correlation.

#### Micro-C

Micro-C libraries were prepared as described (*6*). Briefly, for each replicate library, 2 million cells obtained from visceral adipose tissue were crosslinked with 1% formaldehyde for 10 min at room temperature. Formaldehyde crosslinking was quenched with 0.25 M glycine for 5 min at RT. After washing with 1xPBS, resuspended cells were further crosslinked with 3 mM DSG in PBS for 40 min at RT. Crosslinking was quenched with 0.4 M glycine for 5 min at RT. Cells were resuspended in cold 1x PBS and incubated on ice for 20 min. After collecting cells by centrifugation, cells were washed with buffer MB#1 (10 mM Tris-HCl, pH 7.5, 50 mM NaCl, 5 mM MgCl2, 1 mM CaCl_2_, 0.2% NP-40, 1x Roche cOmplete EDTA-free). Chromatin was fragmented with MNase for 10 min at 37°C and digestion was stopped with 5 mM EGTA at 65°C for 10 min. The chromatin was resuspended in 1x NEBuffer 2.1 (NEB, #B7202S) and dephosphorylated by addition of 5 μl rSAP (NEB, #M0203) at 37°C for 45 min. 5′ overhangs were generated with this pre-mix (50 mM NaCl, 10 mM Tris, 10mM MgCl2, 100 µg/ml BSA, 2 mM ATP, 3 mM DTT 8 μl Large Klenow Fragment (NEB, #M0210L) and 2 μl T4 PNK (NEB, #M0201L)) at 37°C for 15 min. The DNA overhangs were filled with biotinylated nucleotides by addition of 100 μl pre-mix (25 μl 0.4 mM Biotin-dATP (Invitrogen, #19524016), 25 μl 0.4 mM Biotin-dCTP (Invitrogen, #19518018), 2 μl 10 mM dGTP and 10 mM dTTP (stock solutions: NEB, #N0446), 10 μl 10x T4 DNA Ligase Reaction Buffer (NEB #B0202S), 0.5 μl 200x BSA (NEB, #B9000S), 38.5 μl H_2_O) and incubation at 25°C for 45 min. The reaction was stopped by addition of 12 μl 0.5 M EDTA (Invitrogen, #15575038) at 65°C for 20 min. After proximity ligation and removal of unligated ends, the DNA was extracted with phenol/chloroform, purified with DNA Clean & Concentrator Kit (Zymo, #4013), and the library was purified via 1.5% agarose gel electrophoresis. Dynabeads™ MyOne™ Streptavidin C1 beads (Invitrogen, #65001) were added to the sample and incubated at RT for 20 min. The beads were washed twice and resuspended in 50 μl TE buffer. Sequencing libraries were prepared with NEBNext® Ultra II DNA Library Prep Kit for Illumina® (NEB, #E7645) according to the manufacturer’s protocol followed by PCR amplification, sample indexing, and DNA purification. Libraries were paired-end sequenced on an Illumina NovaSeq 6000 instrument. Two biological replicates were generated, and replicates were combined for all analyses after ensuring high correlation.

#### Immunohistochemistry of ARC-ME tissues

Adult 8-16 weeks old mice were anaesthetized with isoflurane and transcardially perfused with ice-cold PBS and 4% PFA. Brains were rapidly extracted and placed into a chilled stainless steel brain matrix with the ventral surface up (catalog no. RBMS-200C, World Precision Instruments, Sarasota, FL). Brains were blocked to obtain a single coronal section containing the hypothalamus (∼ 3mm thick). The hypothalamus tissue was fixed overnight with 4% PFA, washed with PBS, cryoprotected with 30% sucrose, embedded in OCT compound, and then sectioned into 18 μm slices using a Leica cryostat. The sections were washed in PBS, blocked and permeabilized at room temperature with 0.3% Tween-20 and 4% BSA and incubated overnight at 4 °C in blocking solution containing anti-AgRP antibody (R&D Systems; AF634; goat; 1:200 dilution). After washes in PBS, the sections were incubated for 2 h at room temperature with donkey anti-goat Alexa Fluor 488 (Thermo Fisher; Cat. 11055; 1:250 dilution). The sections were washed in PBS, counterstained and mounted on a slide with Vectashield Anti-fade Mounting Medium with DAPI (Cat. H-1200-10, Vector Labs, Burlingame, CA) and sealed with CoverGrip Coverslip Sealant (Cat. 23005, Biotium Inc, Fremont, CA). Fluorescence images were captured with a Zeiss Axio Imager Z1 microscope (Carl Zeiss Imaging), digital camera (Carl Zeiss Imaging, AxioCamHRc) and 20x lens (Plan-Apochromat 20x/0.8), and analyzed with AxioVision SE 4.9.1.

#### Microdissection of hypothalamic ARC-ME tissues and generation snATAC-seq libraries

Mouse brains were dissected after trans cardiac perfusion with cold 1x DPBS, and coronally sliced to obtain 1 mm sections. The ARC-ME region was microdissected in cold 1x DPBS and nuclei were isolated using a pre-chilled Dounce tissue grinder and iodixanol gradient as described previously (*3, 7, 8*) with the addition of 200 units of RNasin Plus Ribonuclease Inhibitor (Promega) in the homogenization buffer. The isolated nuclei were immediately used for single cell MultiOME ATAC + Gene Expression kit according to the manufacturer’s instructions, with the following noted modifications. Briefly, nuclei were resuspended in 1x diluted nuclei buffer (10x Genomics) with 2% BSA (Sigma). We targeted 10,000 nuclei per sample and loaded all samples at 16,100 nuclei. Libraries were barcoded and quantified on a Bioanalyzer (Agilent) before pooling and sequencing. Libraries were sequenced on an Illumina NovaSeq6000 instrument. ATAC libraries were sequenced targeting 25,000 read-pairs per nucleus (R1: 50, R1: 50, i7:8, i5:24) and RNA libraries were sequenced targeting 20,000 read-pairs per nucleus (R1:28, R2: 90, i7: 10, i5:10). Sequencing was performed at the UCSF Center for Advanced Technology (CAT) Genomics core. After sequencing, the matching RNA and ATAC libraries were aligned using Cell Ranger ARC (10x Genomics, v2.0.0) with default parameters, and were mapped against the mm10 mouse genome (redata-cellranger-arc-mm10-2020-A-2.0.0, 10x Genomics). The quality of the snRNA-seq libraries was poor and the RNAs for most genes of interest in our analyses were not present in the nuclear fraction. Therefore, snRNA-seq data were not used.

#### Analysis of ATAC-seq data

Paired reads were aligned to the mouse mm10 reference genome using Bowtie2 (*9*). ATAC-seq reads were aligned using default parameters except -X 2000 -m 1. PCR duplicates were removed using Picard Tools (http://picard.sourceforge.net; https://broadinstitute.github.io/picard/). To adjust for fragment size, we aligned all reads as + strands offset by +4 bp and – strands offset by -5 bp (*10*). For all ATAC-seq datasets subnucleosome size and mono-nucleosome-size reads were separated by choosing fragments 50-115 bp and 180-247 bp in length, respectively. MACS2 (*11*) was used for peak calling of subnucleosomal reads, which represent bound transcription factors.

#### ChIP-seq data processing

All reads were mapped to unique genomic regions using Bowtie2 (*9*) and the mm10 mouse genome. PCR duplicates were removed using Picard Tools (http://picard.sourceforge.net; https://broadinstitute.github.io/picard/). MACS2 (*11*) was used to call peaks using default parameters with IgG ChIP-seq data as a control.

#### RNAseq data processing

The raw RNA-sequencing reads were filtered by FastQC, and all reads were aligned using STAR (v-2.7.1) to the mm10 mouse genome with default parameters. Differentially expressed genes were identified using the R package “Deseq2” with a cut-off *p* value ≤0.05 and fold change ≥2.

#### MeRIP-seq and chromosome-associated RNA m^6^A-seq data analysis

Raw reads were trimmed with Trimmomatic-0.38, then aligned to the mouse genome and transcriptome (mm10) using HISAT2 (version 2.2.0) with ‘--rna-strandness RF’ parameters. Annotation files downloaded from GENCODE database (https://www.gencodegenes.org/) were used. m^6^A peaks were called using MACS with parameter ‘--nomodel’ and ‘--keep-dup 5’.

#### Hi-C and Micro-C data processing

Paired-end reads from Hi-C experiments were aligned to the mouse mm10 reference genome using Juicer (*5*). After PCR duplicates and low-quality reads were removed, high-quality reads were assigned to DpnII restriction fragments, and Hi-C interaction contacts were mapped in a binned matrix to create a hic file. SIP was used to call CTCF loops in the Hi-C interaction matrix (*12*). All statistically significant peaks were post-filtered for observed values greater than 12 Hi-C contacts, observed over expected values greater than 2 Hi-C contacts, and interactions less than 5 Mb apart. The plot connections function in the Cicero package (*13*) was used to obtain arc views of significant loops. Fit-Hi-C (*14*) was used to call significant interaction at 25 kb resolution from the second pass with a q-value threshold of q > 0.001.

#### Processing and analyses of snATAC-seq data

Reads mapping to the mitochondrial genome, chromosome Y, as well as common blacklisted regions were masked and excluded from downstream analysis. ArchR (v1.0.1) (*15*) was used for processing the fragment data, quality control, dimensionality reduction, and clustering with default parameters with the following modifications: 1) Harmony batch correction was performed prior to unsupervised dimensionality reduction, and clustering (*16*); 2) Nuclei were included in the downstream analysis if they had a TSS score ≥4 and more than 1000 unique nuclear fragments; 3) Doublets were removed using the ArchR doublet detection tool using default parameters. Overall, 1303 cells were removed as doublets. After quality control, a total of 19,059 cells were used for downstream analysis with median TSS score of 15.204 and median fragments per nucleus of 16120. The quality control matrix is provided in Supplementary Table 2.Cell type identification was performed based on gene activity scores calculated using ArchR with default parameters; the gene activity scores is correlated with gene expression and calculated based on chromatin accessibility of the gene body, promoter and distal regulatory regions (*7, 15*). Marker genes for each cluster were identified using ArchR’s getMarkerFeatures() function (filtering threshold: FDR < 0.01 & Fold change > 1.4; Supplementary Table 2) and comparison to known marker genes of ARC-ME cell types (*17–22*). Gene score and gene module score were rendered with imputation using MAGIC (*23*) with the addImputeWeights() function in ArchR and utilized on the UMAPs shown in the manuscript. Neuronal clusters were identified based on high gene activity score at the *Tubb3* and *Rbfox3* genes. Sub-clustering was performed on the neuronal and non-neuronal clusters. The ventromedial hypothalamus is located dorsal to the arcuate nucleus. Due to possible variability in the tissues isolated by microdissection, the clusters with high gene module scores for *Sf1*, *Nr5a1*, and *Fezf1* were identified as ventromedial hypothalamus and excluded from further analysis. Excitatory glutamatergic and inhibitory GABAergic neurons were identified based on high gene activity score for the *Slc17a6*, *slc32a1* genes, respectively. AgRP/NPY (cluster 1) and POMC/Cartpt (cluster 13) neurons were identified based on high gene activity score for *Agrp*, *Npy* and *Pomc*, *Cartpt*, respectively. Tanycytes were identified based on high gene activity score at the *Rax* gene. RG-NSCs were identified as previously described (*21*) based on high gene activity score at *Agt, Hes5, Ntsr2, Mfge8, Htra1, Sox2* with low gene score at *Gfap*, a marker of mature astrocytes. NG2-Oligodendrocyte progenitor cells (OPCs) were identified based on high gene activity score at the *Cspg4, Lhfpl3, Gpr17* genes. Mature oligodendrocytes were identified based on high gene activity score at *Mag, Opalin, Mal* (*18*). A detailed list of marker genes for each neuronal and non-neuronal cluster and differential gene activity scores can be found in Supplementary Table 2.

For differential chromatin accessibility analysis of clusters or cell types, non-overlapping pseudobulk replicates for control and BPA libraires were generated from groups of cells. For each cell cluster, the uniform peak matrix from the getPeakSet() was used to calculate the peak coverage per peak in each cell cluster using the getGroupSE() function. These pseudobulk replicates were then used for differential peak comparisons using DESeq2 (v.1.32.0), and only peaks with FDR<0.05 and log2(fold change) > 1 were considered differential (*24*). GetGroupBW() was used to generate pseudobulk ATAC-seq signal track for each cell cluster per replicate or combined for treatments, and visualized using the Integrative Genomics Viewer (IGV).

## Acknowledgements

We would like to thank Dr. Cynthia Vied at the Translational Science Laboratory of Florida State University and the Center for Advanced Technologies (CAT) sequencing core at UCSF for help with Illumina Sequencing; Drs. Karolina Piotrowska-Nitsche and Christopher Raymond from the Emory Mouse Transgenic and Gene Targeting Core for their assistance with the embryo transfer; and Dr. Fred Strobel from the Emory Mass Spectrometry Core for his assistance with detection of BPA in mouse embryos. This work was supported by U.S. Public Health Service Awards R01 ES027859 and P30ES019776 (VGC); R35 NS111602, R01 HG008935, U01 MH116441 (PJ); and R00AG059918 (MRC) from the National Institutes of Health. H-LW was supported by NIH F32 ES031827. BJB was supported by NIH T32 GM008490. The content is solely the responsibility of the authors and does not necessarily represent the official views of the National Institutes of Health.

## Author Contributions

YHJ planned and performed experiments, analyzed data, and contributed to writing the manuscript; H-LW performed experiments and analyzed data, and contributed to writing the manuscript; HL performed experiments; BJB performed experiments and analyzed data; AMS performed experiments; J-FX performed experiments; DR performed experiments and analyzed data; FG performed experiments and analyzed data; PJ planned experiments; MRC planned experiments; VGC planned experiments and wrote the manuscript.

## Data Availability

ChIP-seq, ATAC-seq, RNA-seq, m^6^A, Hi-C, micro-C and snATAC-seq data are available from NCBI’s Gene Expression Omnibus (GEO). The accession number for all the datasets reported in this paper is GSE149309. Reviewers can access these data using token mjwvqswoljmbziz.

## Code Availability

Custom scripts were used to separate ATAC-seq reads into subnucleosomal and nucleosome-size ranges. These scripts are available without restrictions upon request.

## Supplemental Figures and Tables

**Supplemental Figure S1.**
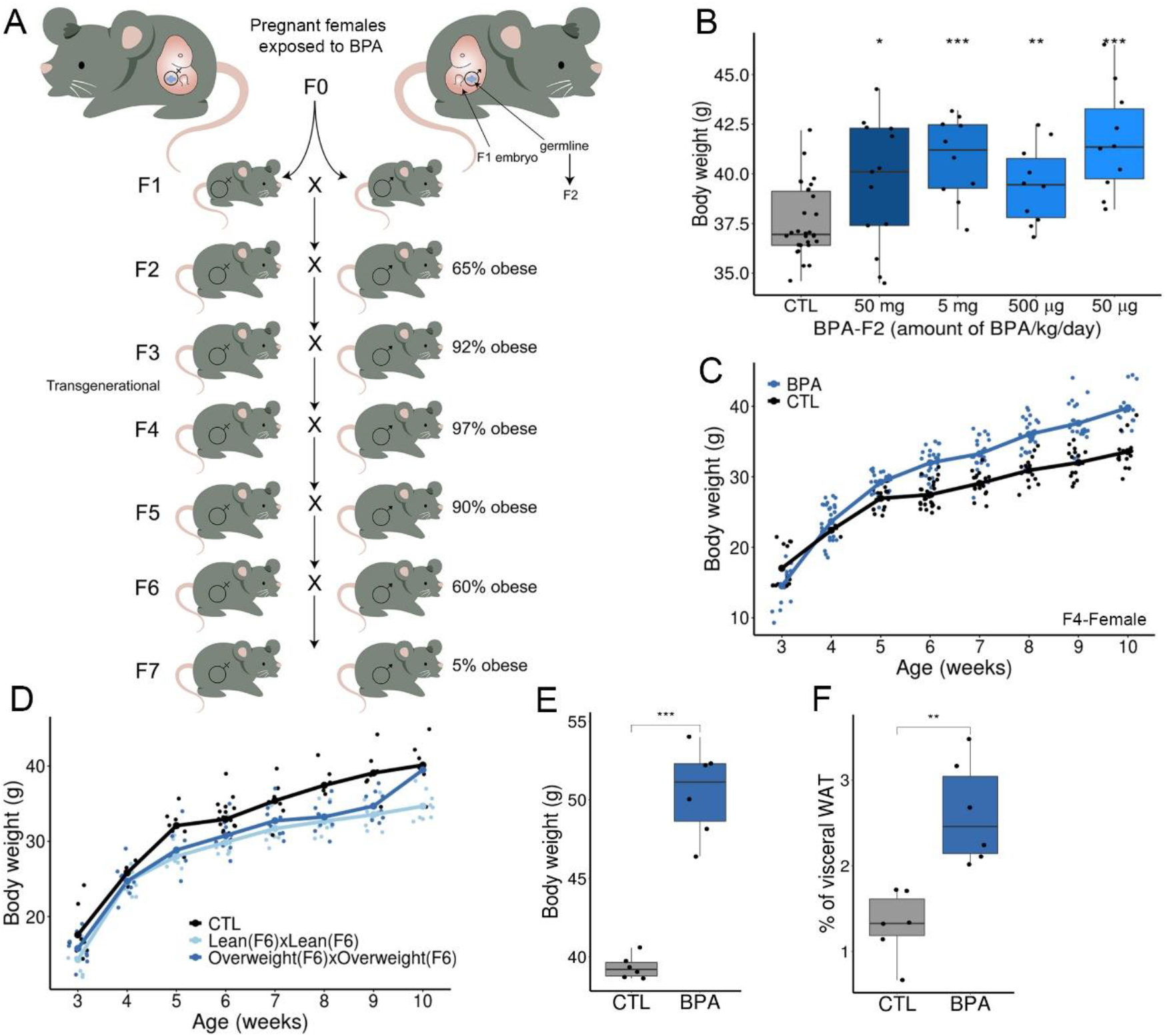
Exposure of pregnant females to BPA causes obesity that can be transmitted transgenerationally. **(A)** Two different pregnant CD1 females (F0) were injected with BPA during days E7.5-E13.5 of embryonic development when the germline of the embryo (labeled in blue) is undergoing demethylation. Male or female F1 progeny from each F0 female are crossed to obtain the F2 generation, which arises from the BPA-exposed germline. F2 males and females are crossed to obtain F3, which is the first generation whose cells were not directly exposed to BPA. The percentage of obese mice in each generation is indicated. **(B)** Exposure to different amounts of BPA results in obesity (CTL n=30, 50 mg n=13, 5 mg n=10, 500 µg n=10, 50 µg n=10). **(C)** Changes in body weight with time for BPA-F4 females (CTL n=16, BPA n=17). **(D)** Changes in body weight with time for control and F7 males arising from crosses between BPA-F6 lean or BPA-F6 overweight animals (CTL n=7, Lean(F6) x Lean (F6) n=8, Overweight (F6) x Overweight (F6) n=9). **(E)** Distribution of body weights for BPA-F4 males (CTL n=6, BPA n=6). **(F)** Distribution of ratios of visceral fat with respect to body weight for BPA-F4 males (CTL n=6, BPA n=6).

**Supplemental Figure S2.**
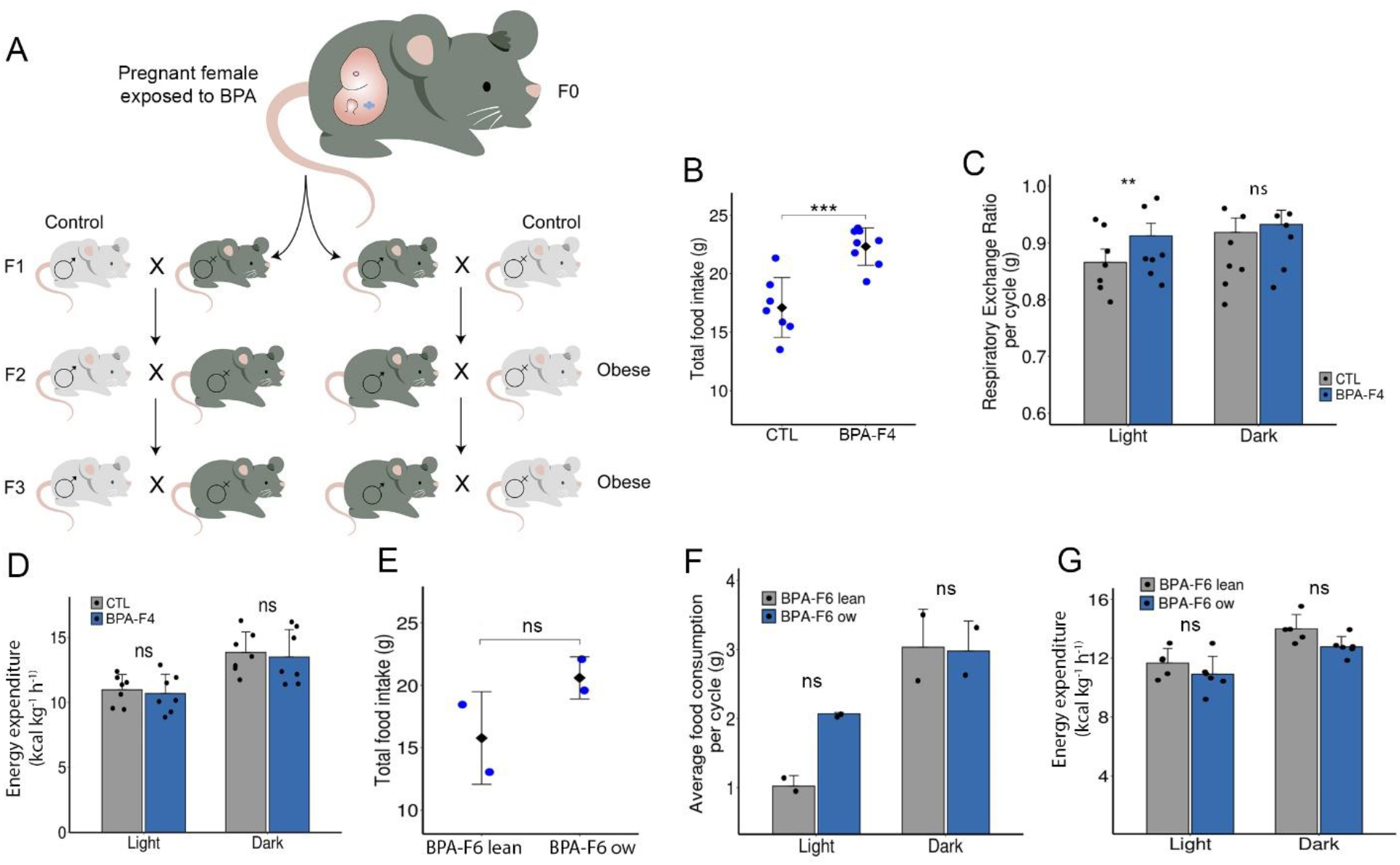
Transgenerational effects of BPA on body weight. (**A**) Outcrosses between BPA-F1 males or females and control mice of the opposite sex. **(B)** Differences in total daily food intake between control and BPA-F4 males (CTL n=7, BPA n=8). **(C)** Respiratory exchange ratio during the light and dark cycles in control and BPA-F4 males (CTL n=7, BPA n=7). **(D)** Energy expenditure during the light and dark cycles in control and BPA-F4 males (CTL n=7, BPA n=7). **(E)** Differences in total daily food intake between overweight and lean BPA-F6 males (BPA-F6 lean n=2, ow n=2). **(F)** Food consumption during the light and dark cycles in control and BPA-F6 overweight and lean males (BPA-F6 lean n=2, ow n=2). **(G)** Energy expenditure during the light and dark cycles in BPA-F6 overweight and lean males (BPA-F6 lean n=5, ow n=6).

**Supplemental Figure S3.**
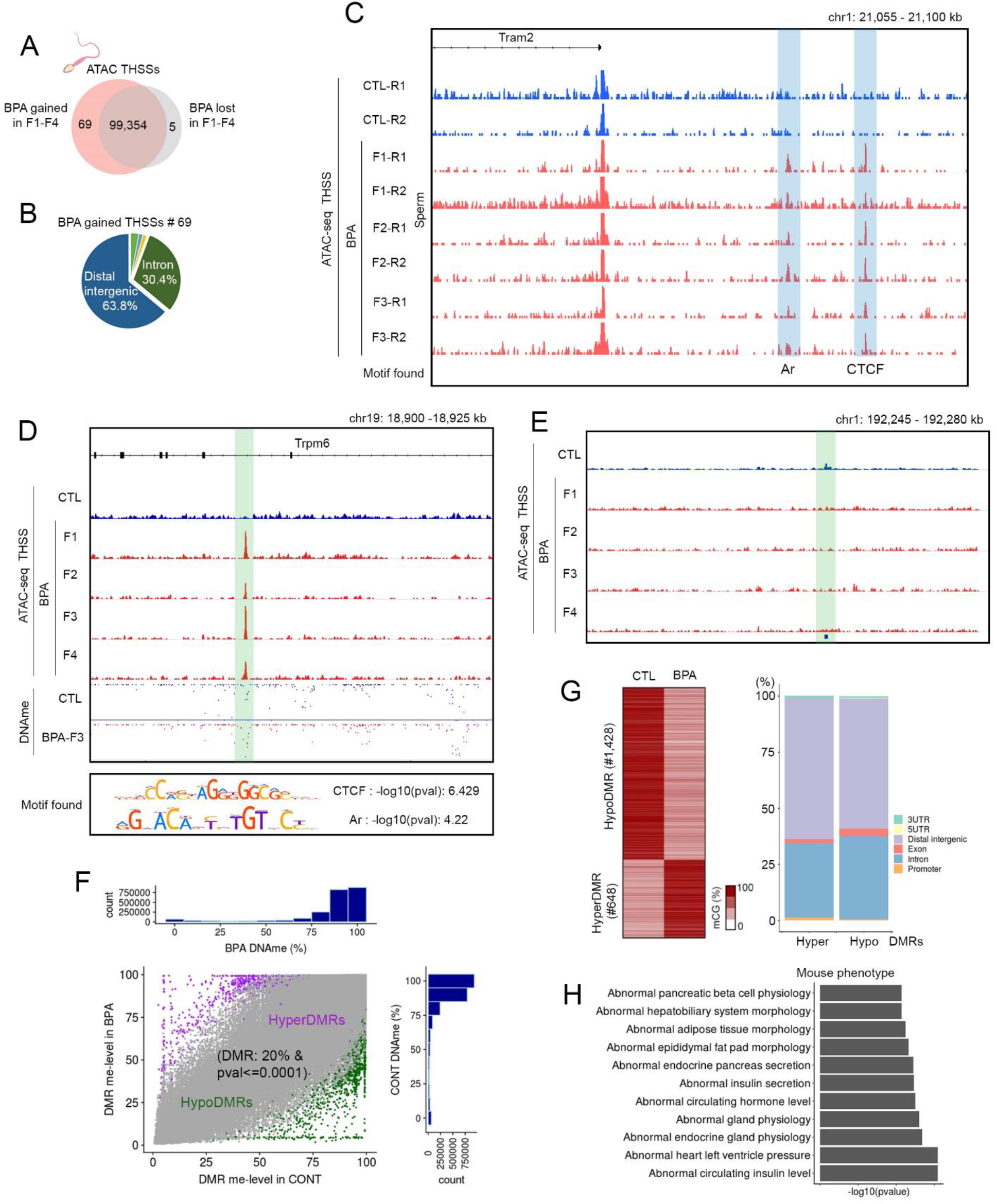
New ATAC-seq sites in BPA-Fi are differentially methylated. **(A)** Overlap between ATAC-seq peaks in sperm for BPA-F1-F4 and controls showing peaks gained or lost in the progeny of BPA-exposed females. **(B)** Distribution of ATAC-seq peaks present in BPA-F1-F4 but not in controls with respect to gene features. **(C)** An example of an ATAC-seq peak present in control but not in BPA-F1-F3. **(D)** An example of an ATAC-seq peak present in BPA-F1-F4 but not in controls in the *Trpm6* gene. The summit of the peak contains motifs for CTCF and Ar. The site is hypomethylated in BPA-F3 with respect to control. **(E)** Example of an ATAC-seq peak lost in BPA-F3 mice with respect to control. **(F)** Analysis of GWBS-seq data showing differentially methylated regions (DMRs) between BPA-F3 and control sperm. **(G)** Heatmap showing different DNA methylation levels at regions hyper- and hypo-methylated in BPA-F3 sperm with respect to control (left). DMRs are enriched at intergenic and intronic regions, suggesting they may correspond to enhancer sequences (right). **(H)** Phenotypes of mice mutant in genes contacted by the 69 ATAC-seq sites present in BPA-F1-F4 but not in controls.

**Supplemental Figure S4.**
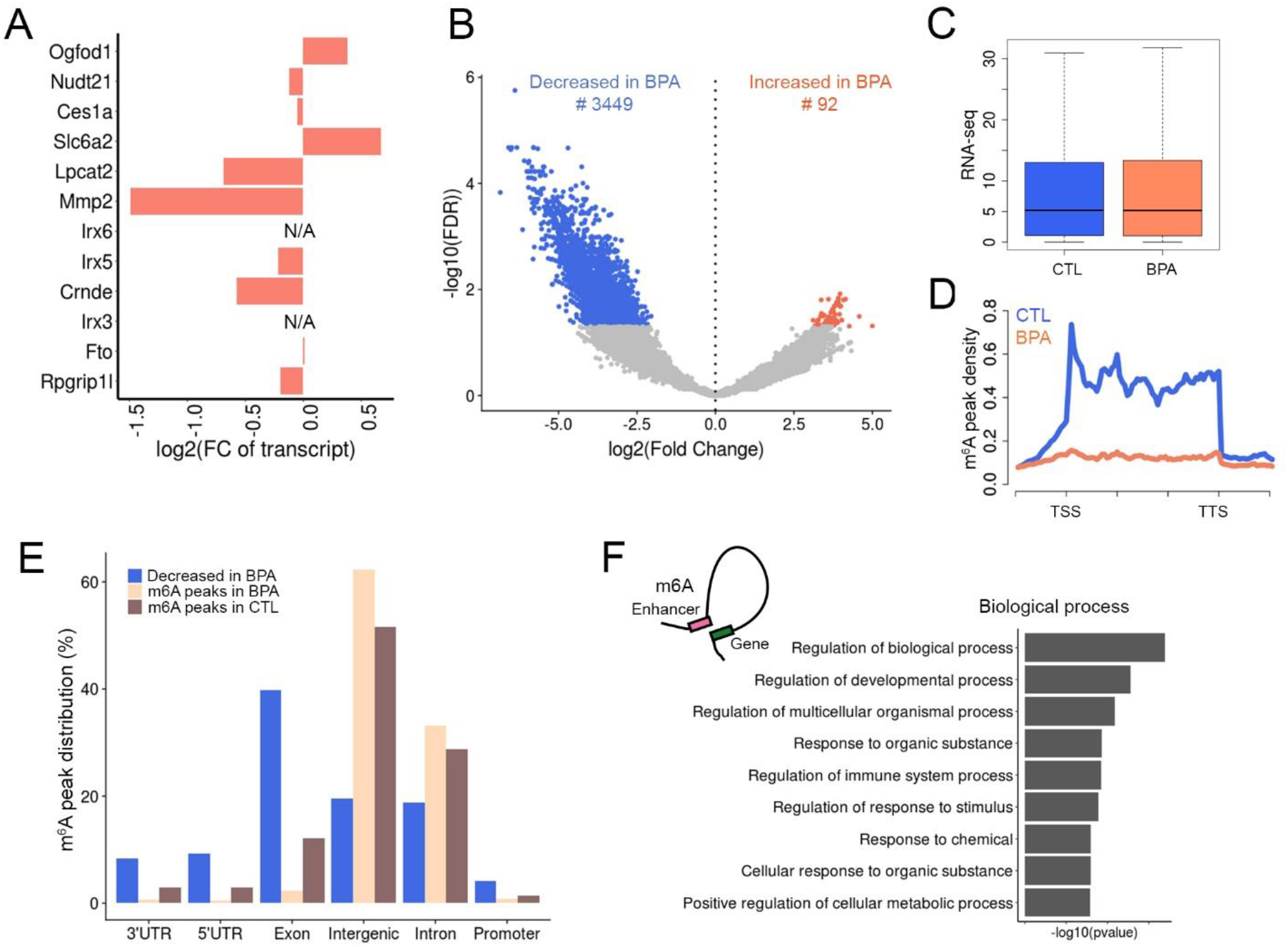
Effect of BPA-induced epigenetic alterations in *Fto* on m^6^A levels in sperm RNAs. **(A)** relative RNA levels in sperm of BAP-F3 and control transcribed from genes present in the region surrounding *Fto*. **(B)** RNAs whose m^6^A levels are altered in BPA-F3 sperm with respect to control. **(C)** Global RNA levels are not different between sperm of BPA-F3 and control mice. **(D)** Levels of m^6^A at sites located in mRNAs are lower in sperm from BPA-F3 mice with respect to control. **(E)** Location of sites in RNAs modified by m^6^A with respect to genomic features. **(F)** Biological processes involving genes whose promoters are contacted by enhancers whose eRNAs show decreased levels of m^6^A in BPA-F3 with respect to control.

**Supplemental Figure S5.**
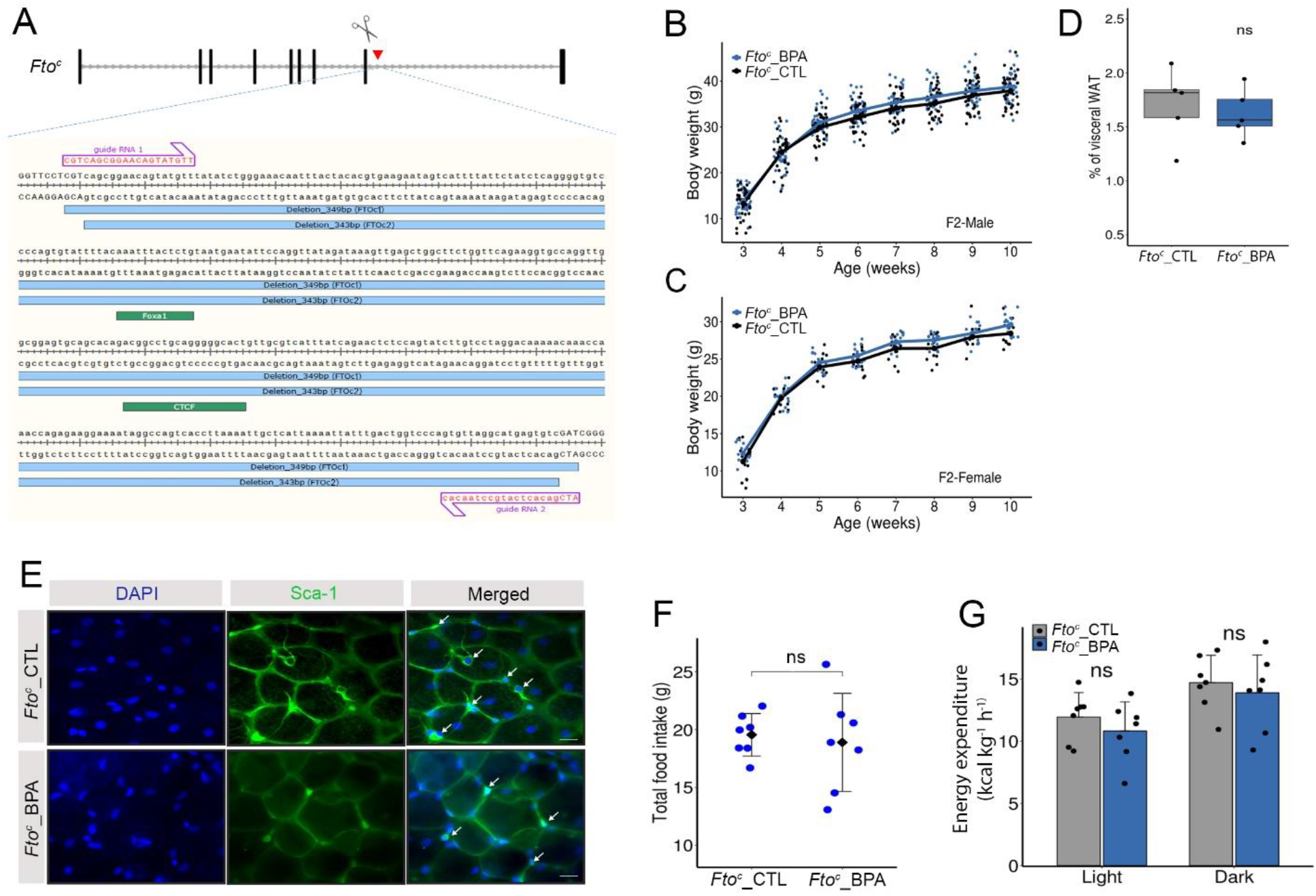
Response of mice carrying the *Fto^c^* allele to BPA exposure. **(A)** Region of intron 8 containing the FOXA1 and CTCF sites deleted in *Fto^c^* mice. Two slightly different deletions gave the same results in all analyses. **(B)**. Weight gain with age in F2 *Fto^c^* male mice ancestrally exposed to BPA (*Fto^c^*_CTL n=35, *Fto^c^*_BPA n=39). **(C)** Weight gain with age in F2 *Fto^c^* female mice ancestrally exposed to BPA (*Fto^c^*_CTL n=11, *Fto^c^*_BPA n=15). **(D)** Percentage of visceral fat with respect to body weight in *Fto^c^* male mice ancestrally exposed or non-exposed to BPA (*Fto^c^*_CTL n=6, *Fto^c^*_BPA n=6). **(E)** Immunofluorescence microscopy of visceral fat tissue from exposed and unexposed *Fto^c^* mice stained with antibodies to Sca-1, which labels adipocyte progenitor cells. **(F)** Total food intake by *Fto^c^* mice ancestrally exposed to BPA and unexposed controls (*Fto^c^*_CTL n=7, *Fto^c^*_BPA n=7). **(G)** Average energy expenditure during the light and dark light cycles by *Fto^c^* mice ancestrally exposed to BPA and unexposed controls (*Fto^c^*_CTL n=7, *Fto^c^*_BPA n=7).

**Supplemental Figure S6.**
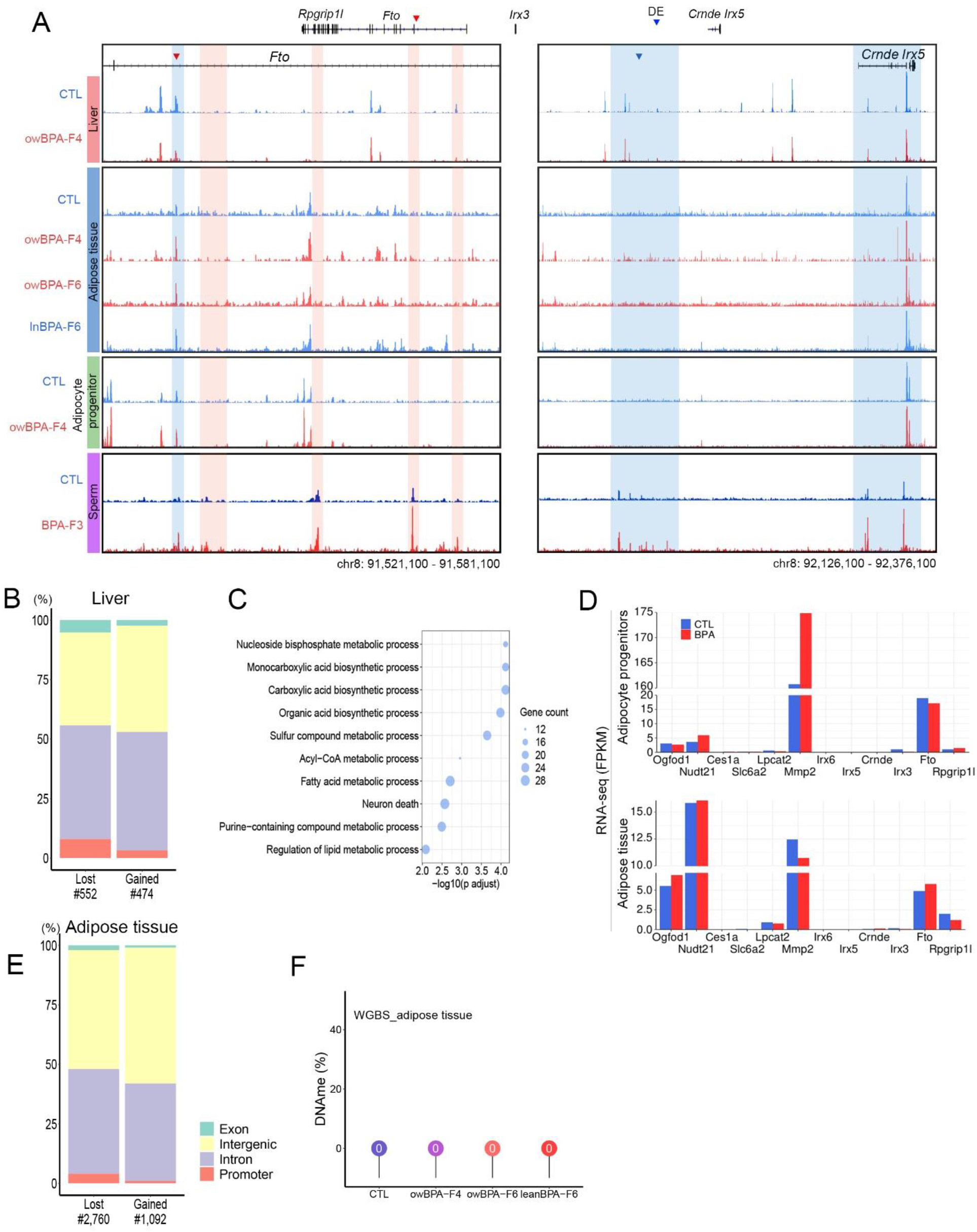
Changes in ATAC-seq and DNA methylation in adult tissues of control and BPA-F4 mice. **(A)** Genome browser view of intron 8 of the *Fto* gene, which contains the proximal enhancer, and the region surrounding *Irx5*, which contains the distal enhancer, showing ATAC-seq peaks in various adult tissues of control and BPA-F4 males. **(B)** Genomic distribution of differential ATAC-seq peaks from liver tissue of control and BPA-F4 males. **(C)** Processes from Go analyses involving genes adjacent to differential ATAC-seq peaks in liver of control and BPA-F4 mice. **(D)** Gene expression for the specified genes from RNA-seq in adult tissues from control and BPA-F4 males. **(E)** Genomic distribution of differential ATAC-seq peaks from visceral fat tissue of control and BPA-F4 males. **(F)**. Changes in DNA methylation at the C at position 2 of the CTCF motif located in the proximal *Fto* enhancer in visceral fat tissue of unexposed control, BPA-F4, overweight BPA-F6, and lean BPA-F6 males at differential ATAC-seq peaks.

**Supplemental Figure S7.**
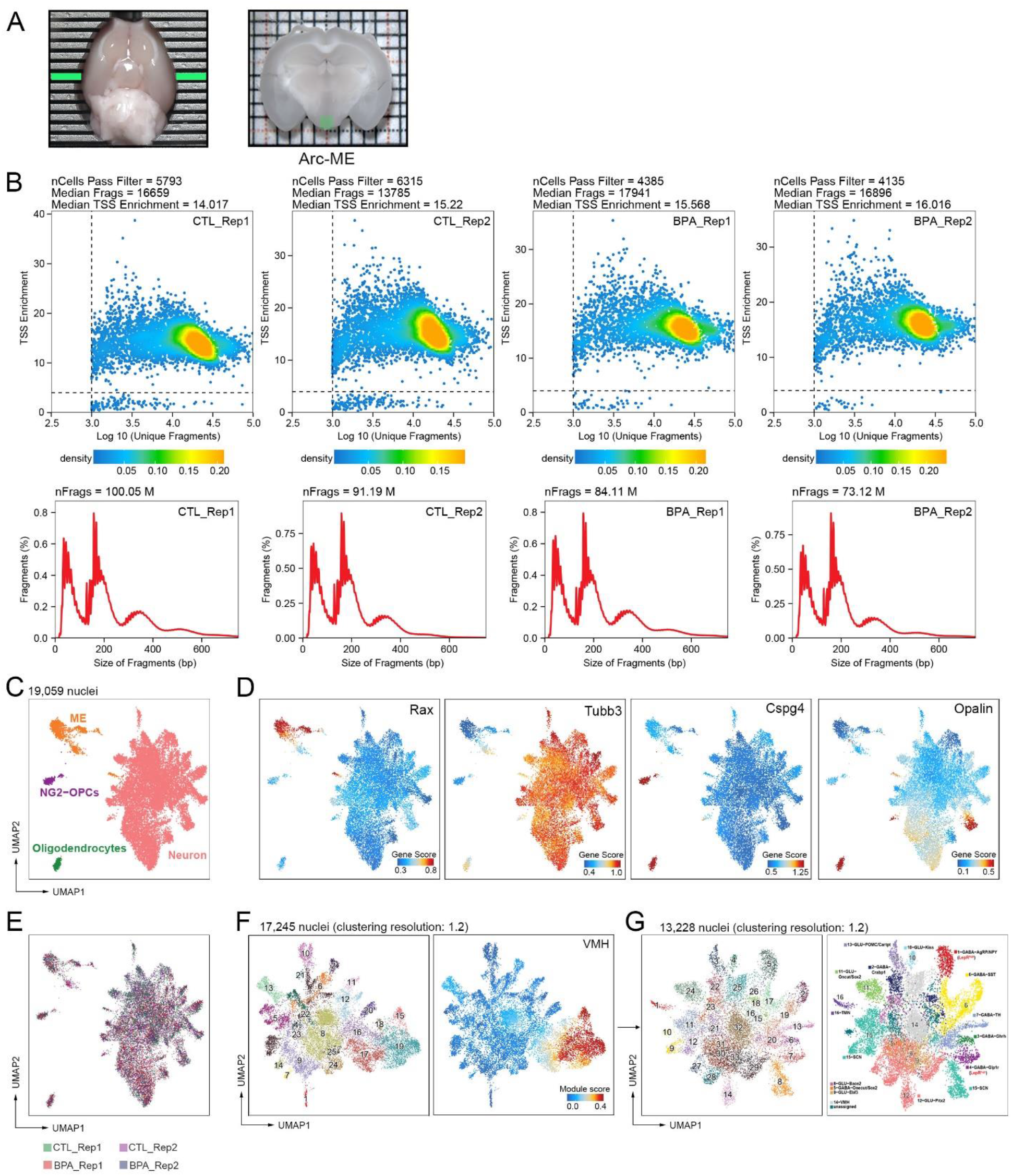
Characterizations of ARC-ME and proximal region using snATAC-seq. **(A)** Mouse brain showing the region dissected in a 1 mm coronal slice using a brain matrix (left panel). Green shaded area marks the ARC-ME region dissected from the coronal slice for snATAC-seq experiments (left panel). **(B)** Strict quality controls of snATAC-seq applied to remove low-quality cells. Top row, 2D-density plots show the unique nuclear fragments and TSS enrichment score per single nucleus for each snATAC-seq library. Nuclei with ≥4 TSS enrichment score and >1000 unique nuclear fragments were considered good quality and used for unsupervised clustering. Bottom row shows the fragment size distribution for each snATAC-seq library corresponding to nucleosome periodicity. **(C)** UMAP projection of 19,059 high quality nuclei colored by major cell types **(D)** UMAP plots showing gene activity score of cell-type and cluster-specific markers. *Rax* is a tanycyte marker, *Tubb3* is a neuronal marker, *Cspg4* is a NG2-OPC and oligodendrocyte marker, and *Opalin* is a mature oligodendrocyte marker. **(E)** UMAP plot implemented with Harmony batch correction of nuclei from CTL and BPA samples, colored by replicate (n= 5,261 and 6,009 for CTLs, n= 4,037 and 3,752 for BPA; related to panel C-D). **(F)** UMAP projections of subclustering analysis of 17,245 neurons (left) and UMAP plot showing the gene module score of the ventromedial hypothalamus (VMH) region (right). **(G)** UMAP projections from subclustering analysis of 13,228 non-VMH neurons (left) and grouped and colored by known ARC-ME neurons and proximal regions. The cell type specific gene activity scores are shown in Supplemental Figure S8.

**Supplemental Figure S8.**
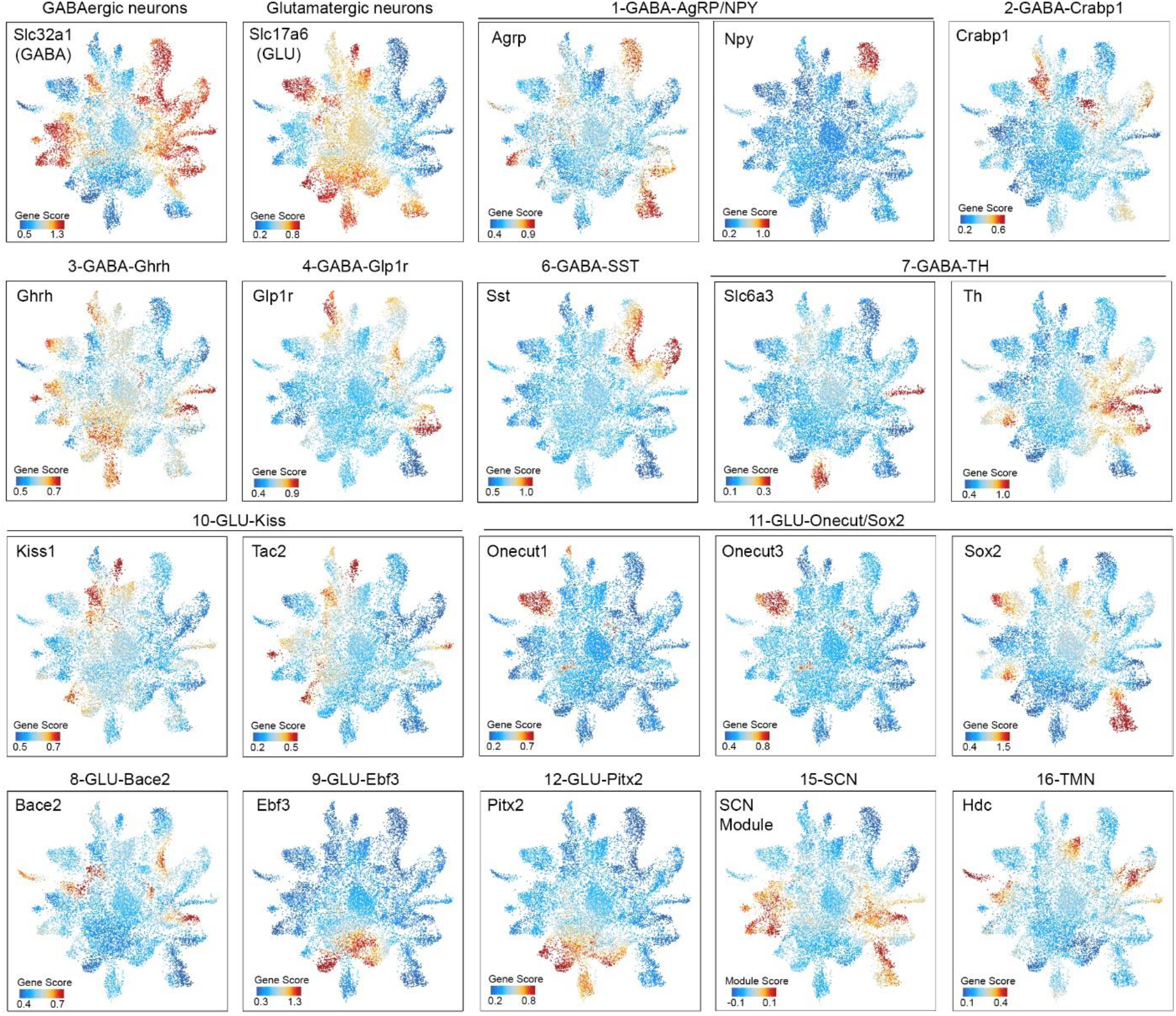
Gene activity score of neuron-type specific markers in snATAC-seq data. A total of 13,228 neurons are shown. SCN module score calculated using a group of SCN marker genes (*Cck, Rgs16, Per1, Per2, C1ql3, Dlx1, Dlx2*) is also shown.

**Supplemental Figure S9.**
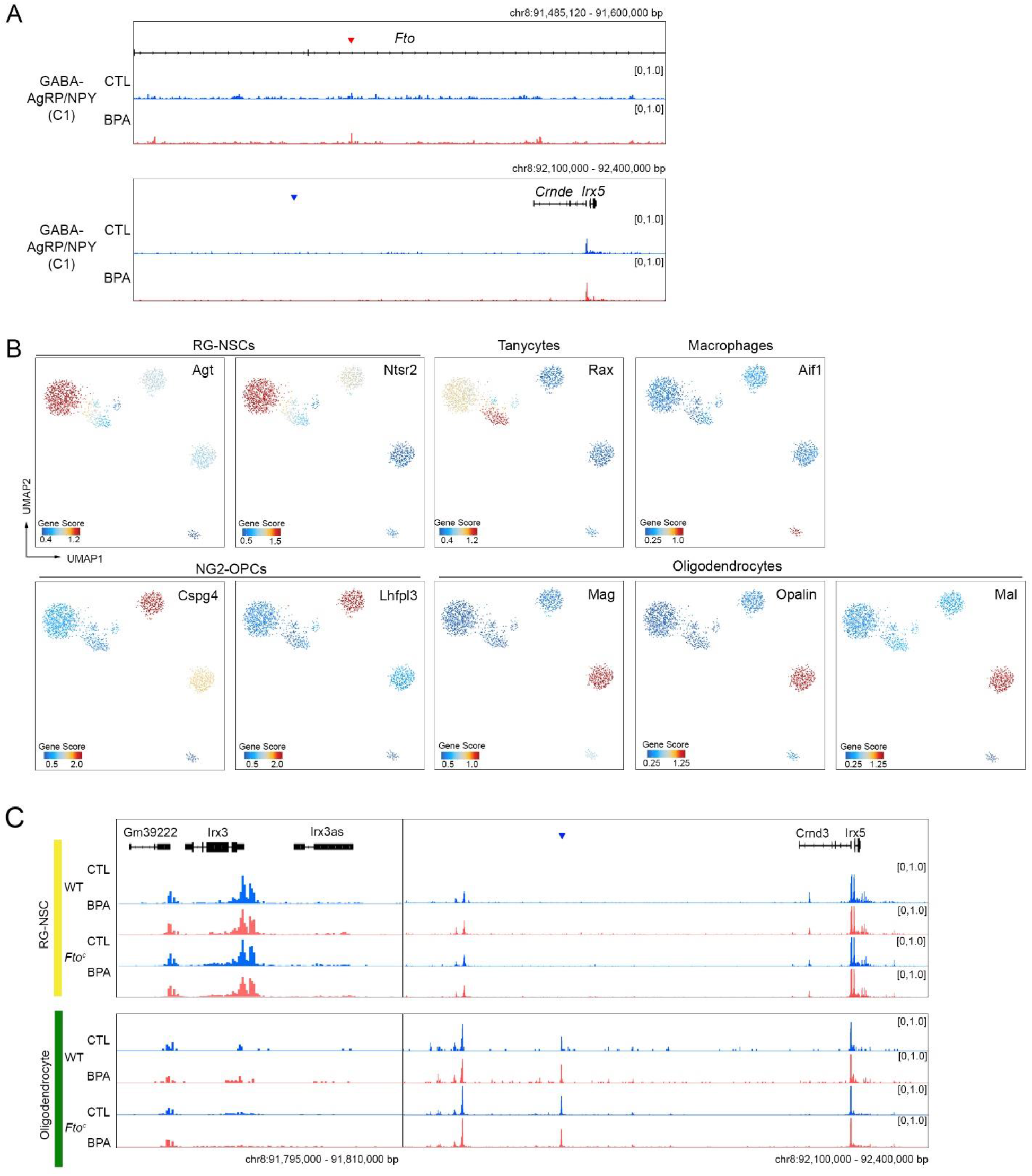
**(A) Chromatin accessibility and clustering analyses of neuronal and non-neuronal cells.** Chromatin accessibility in the regions surrounding the *Fto* proximal and distal enhancers. Pseudo-bulk chromatin accessibility in AgRP/NPY (cluster 1) and POMC/Cartpt (cluster 13) neurons in snATAC-seq datasets from WT unexposed and BPA exposed mice. **(B)** UMAP projections and cell type specific gene activity score from subclustering analysis of 1,527 non-neuronal nuclei. **(C)** Chromatin accessibility in the regions surrounding the *Fto* proximal and distal enhancers. Pseudo-bulk chromatin accessibility in RG-NSCs and oligodendrocytes in snATAC-seq datasets from WT unexposed and BPA exposed mice.

**Supplemental Figure S10.**
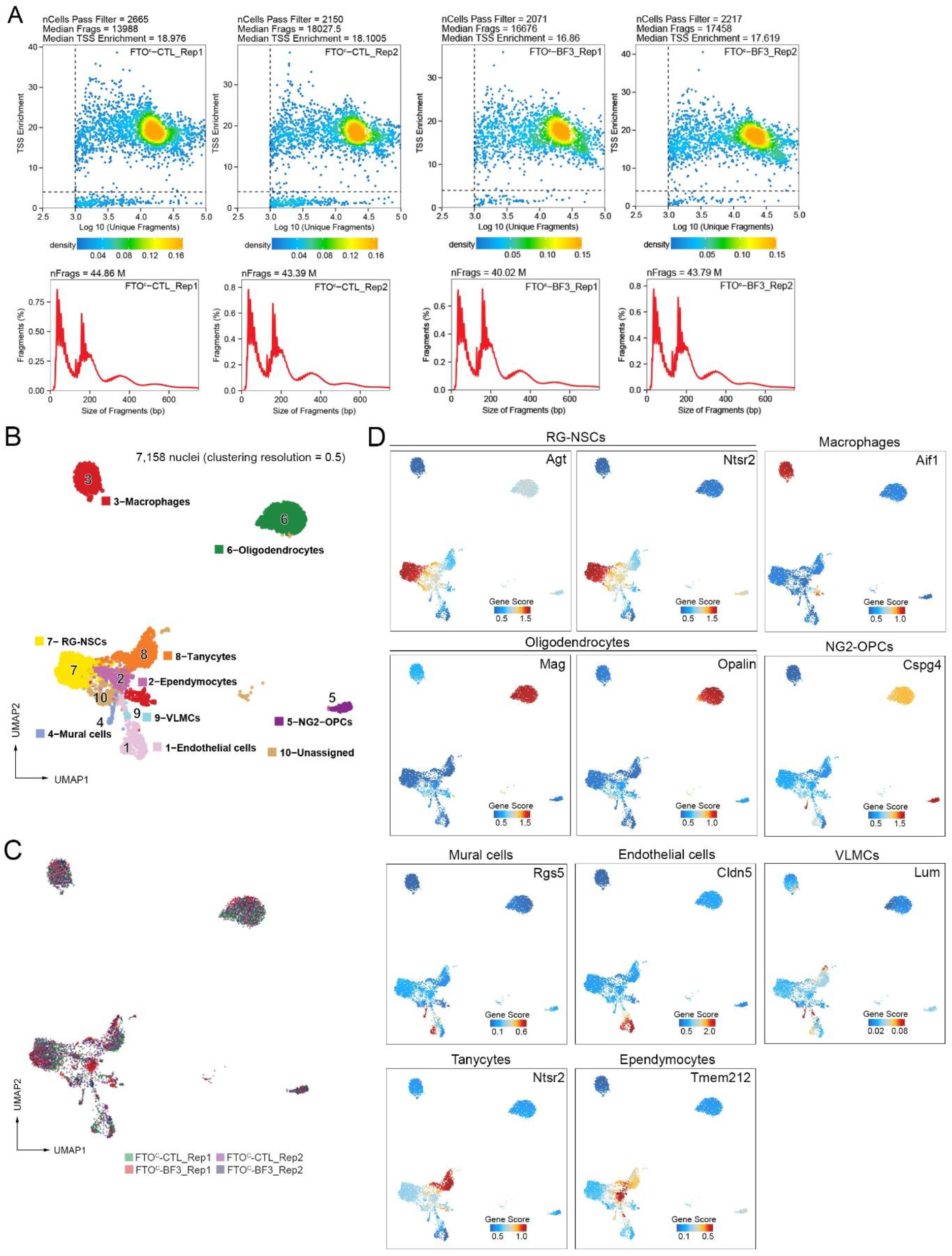
snATAC-seq analysis of *Fto^c^* mice ancestrally exposed to BPA and unexposed controls. **(A)** Quality control analysis of snATAC-seq dataset obtained in *Fto^c^* mice. Top row, 2D-density plots show unique nuclear fragments and TSS enrichment scores per single nucleus for each snATAC-seq library. Nuclei with ≥4 TSS enrichment score and >1000 unique nuclear fragments were considered good quality and used for unsupervised clustering. Bottom row shows the fragment size distributions for each snATAC-seq library corresponding to nucleosome periodicity. **(B)** UMAP projections of subclustering analysis of 7,158 none-neuronal nuclei. **(C)** UMAP plot implemented with Harmony batch correction of nuclei from *Fto^c^*-CTL and *Fto^c^*-BPA mice, colored by replicate (n= 2,208 and 1,735 for *Fto^c^*-CTLs, n= 1,568 and 1,647 for *Fto^c^*-BPA). **(D)** Cell type specific gene activity scores for various non-neural markers.

## Supplemental Tables

Supplemental Table 1 and 2 are provided as Excel files.

**Table S1.** Quality control and mapping statistics of Hi-C and Micro-C libraries.

**Table S2.** Clustering, cell type annotation and summary quality control statistics of scATAC-seq

